# NLRP3 inflammasome contributes to host defense against *Talaromyces marneffei* infection

**DOI:** 10.1101/2020.08.17.253518

**Authors:** Haiyan Ma, Jasper FW Chan, Yen Pei Tan, Lin Kui, Chi-Ching Tsang, Steven LC Pei, Yu-Lung Lau, Patrick CY Woo, Pamela P Lee

**Affiliations:** Department of Pediatrics and Adolescent Medicine, Li Ka Shing Faculty of Medicine, The University of Hong Kong, Hong Kong SAR, China; Department of Microbiology, Li Ka Shing Faculty of Medicine, The University of Hong Kong, Hong Kong SAR, China

## Abstract

*Talaromyces marneffei* is an important thermally dimorphic pathogen causing disseminated mycoses in immunocompromised individuals in southeast Asia. Previous study has suggested that NLRP3 inflammasome plays a critical role in antifungal immunity. However, the mechanism underlying the role of NLRP3 inflammasome activation in host defense against *T. marneffei* remains unclear. We show that *T. marneffei* yeasts but not conidia induce potent IL-1β response, which is differentially regulated in discrete immune cell types. Dectin-1/Syk signaling pathway mediates pro-IL-1β production, and NLRP3 inflammasome is activated to trigger the processing of pro-IL-1β into IL-1β. The activated NLRP3 inflammasome partially promotes Th1 and Th17 immune responses against *T. marneffei* yeasts. *In vivo*, mice with NLRP3 or caspase-1 deficiency exhibit higher mortality rate and fungal load compared to wild-type mice. Herein, our study provides the first evidence that NLRP3 inflammasome contributes to host defense against *T. marneffei* infection, which may have implications for future antifungal therapeutic designs.

## Introduction

*Talaromyces marneffei* is a dimorphic fungus geographically restricted to Southeast Asia. It causes a form of endemic mycosis that predominantly affects individuals with HIV infection and ranks the third most common opportunistic infection in acquired immunodeficiency syndrome (AIDS), following tuberculosis and cryptococcosis (Sirisanthana & Supparatpinyo, 1998; Supparatpinyo *et al*, 1994; Vanittanakom *et al*, 2006; Cao *et al*, 2019). In highly endemic regions such as Chiang Mai in Thailand and Haiphong in Vietnam, *T. marneffei* is in fact the most common systemic opportunistic fungal infection in AIDS (Vanittanakom *et al,* 2006; Son *et al*, 2014). Compared with HIV-negative individuals, fungemia, hepatomegaly and splenomegaly occur much more commonly in HIV-positive patients, supporting profound T-lymphopenia as the key predisposing factor for disseminated talaromycosis (Kawila *et al*, 2013).

Less commonly, talaromycosis occurs in other immunocompromised conditions such as solid organ or hematopoietic stem cell transplantation, autoimmune diseases or primary immunodeficiency diseases (PID) (Chan *et al*, 2016; Lee *et al*, 2012; Lee *et al,* 2014; Lee & Lau, 2017; Luo *et al*, 2010; Stathakis *et al*, 2015). *T. marneffei* shares some common characteristics with other phylogenetically related dimorphic fungi such as *Histoplasma* and *Coccidioides.* They exist naturally in the soil environment as conidia which enter the human body via inhalation route, and are converted to the yeast form after being phagocytosed by tissue-resident macrophages. These dimorphic fungal pathogens commonly result in systemic infection when immunity is suppressed, and the uncontrolled proliferation of yeasts in the macrophages eventually lead to dissemination via the reticuloendothelial system. *Paracoccidioides brasiliensis* and *P. lutzii*, another dimorphic fungi which is endemic in Latin America, is prevalent in immunocompetent individuals and co-infection with HIV is uncommon, whereas talaromycosis, histoplasmosis and coccidioidomycosis are regarded as AIDS-defining conditions in their respective endemic regions (Limper *et al*, 2017). They have also been reported in patients with autoantibodies against interferon gamma (anti-IFNγ), and PID characterized by impaired Th17 immune response such as autosomal dominant (AD) hyper-IgE syndrome caused by loss-of-function STAT3 defect and AD chronic mucocutaneous candidiasis (CMC) caused by gain-of-function STAT1 defect, as well as CD40L deficiency and IFN-γ receptor deficiency. These acquired defects and inborn errors of immunity provide evidence on the critical role of Th1 and Th17 immune response in host defense against these dimorphic fungi (Lee & Lau, 2017; Lee *et al*, 2019). However, knowledge about host-pathogen interaction in endemic mycoses is much less comprehensive than other common opportunistic fungal pathogens of worldwide distribution. In particular, little is known about how *T. marneffei* is recognized by innate immune cells and the ensuing cytokine signaling that triggers adaptive immune response.

Immune responses against fungal infection are mounted by fungal pathogen-associated molecular patterns (PAMPs) that can be recognized by pathogen recognition receptors (PRRs), including Toll-like receptors (TLRs), C-type lectin receptors (CLRs), NOD-like receptors (NLRs) (Erwig & Gow, 2016). CLRs are central for fungal recognition, uptake and induction of innate and adaptive immune responses (Vautier *et al*, 2012). Dectin-1 specifically recognizes β-1,3-glucans in fungal cell wall (Taylor *et al*, 2007), while Dectin-2 and macrophage inducible C-type lectin (Mincle) recognize α-mannan and α-mannose, respectively (Vautier *et al*, 2012). Dectin-1, Dectin-2 and Mincle signal through the Syk-CARD9 pathway which induces NF-kB activation as well as cytokine and chemokine gene expression (Mócsai *et al*, 2010). NLRP3 is a member of NLRs family, which is able to indirectly sense danger signals, e.g. PAMPs (Schroder & Tschopp, 2010). Upon recognition, NLRP3 interacts with adaptor protein ASC that can further recruit pro-caspase-1, leading to the assembly of multiprotein complex known as NLRP3 inflammasome (Latz *et al*, 2013). Thereafter, pro-caspase-1 can be activated through auto-proteolysis, resulting in the release of caspase-1 p10 and p20 (Schroder & Tschopp, 2010). Active caspase-1 not only processes pro-IL-1β and pro-IL-18 into bioactive IL-1β and IL-18, but also mediates pyroptosis (Man & Kanneganti, 2016). The activation of IL-1β transcription and NLRP3 inflammasome is dependent on the Dectin-1/Syk signaling pathway in human macrophages (Kankkunen *et al*, 2010). The activation of Dectin-1 also triggers the induction of IL-23 through the Syk-CARD9 signaling pathway and drives robust Th17 response (LeibundGut-Landmann *et al*, 2007). The IL-17/IL-23 axis has been shown to promote immunity to *C. albicans*, *Aspergillus* and dimorphic fungi such as *H. capsulatum* (Deepe Jr. & Gibbons, 2009; Chamilos *et al*, 2010; Wu *et al*, 2013) and *Paracoccidioides brasiliensis* (Tristão *et al*, 2017). In a murine model of disseminated candidiasis, NLRP3 inflammasome exerts antifungal activity through driving protective Th1 and Th17 immune responses (van de Veerdonk *et al*, 2011). NLRP3 inflammasome has also been shown to mediate IL-1β and IL-18 signaling in murine DC and macrophages infected by *P. brasiliensis* (Tavares *et al*, 2013; Feriotti *et al*, 2015; Ketelut-Carneiro *et al,* 2015), and is essential for the induction of protective Th1/Th17 immunity against pulmonary paracoccidioidomycosis (Feriotti *et al*, 2017; de Castro *et al*, 2018).

Previous study showed that IL-1β gene transcription is upregulated in human monocyte-derived DC and macrophages stimulated by *T. marneffei* (Ngaosuwankul *et al*, 2008). However, the mechanism of IL-1β induction has not been explored, and whether NLRP3 inflammasome plays a role in protective immunity against *T. marneffei* is unknown. From our earlier work, impaired Th1 and Th17 effector functions emerges as the common pathway in various types of PIDs rendering susceptibility to talaromycosis (Lee *et al*, 2012; Lee *et al*, 2014; Lee *et al*, 2019; Lee & Lau, 2017). In this study, we sought to investigate the role of NLRP3 inflammasome in the induction of Th1 and Th17 immune response to *T. marneffei*. Our findings show that Dectin-1/Syk signaling pathway mediates pro-IL-1β production, and NLRP3 inflammasome is assembled to enhance the processing of pro-IL-1β and pro-IL-18 into mature IL-1β and IL-18 in response to *T. marneffei* yeasts. *T. marneffei* induces Th1 and Th17 cell differentiation, which is attenuated by caspase-1 inhibition. Furthermore, our *in vivo* studies show that NLRP3 and caspase-1 confer protection against disseminated *T. marneffei* infection.

## Results

### *T. marneffei* yeasts, but not conidia, induce potent IL-1β response in human PBMCs

To dissect the production of cytokines elicited by *T. marneffei*, freshly isolated human peripheral blood mononuclear cells (PBMCs) were co-cultured with *T. marneffei* yeasts or conidia at 0.5 multiplicity of infection (MOI) for indicated incubation time points. As shown in Fig 1A, IL-1β became detectable at 8 hr, peaked at 18 hr, and remained plateaued thereafter till day 5 post infection in the culture supernatant of human PBMCs infected with live *T. marneffei* yeasts. On the other hand, TNF-α production was detectable as early as 4 hr, and rapidly peaked at 18 hr, and subsequently showed a slight falling trend up to day 5 post infection (Fig 1B). These data demonstrated that *T. marneffei* yeasts were a potent inducer of IL-1β and TNF-a production. In contrast to *T. marneffei* yeasts, only minimal levels of IL-1β was detected in PBMCs co-cultured with conidia at 5 days post infection (Fig 1C), whereas TNF-α production began to increase steadily after 2 days of incubation (Fig 1D). This suggests that *T. marneffei* conidia were able to trigger TNF-α production in PBMCs but failed to induce IL-1β response.

**Figure 1.**
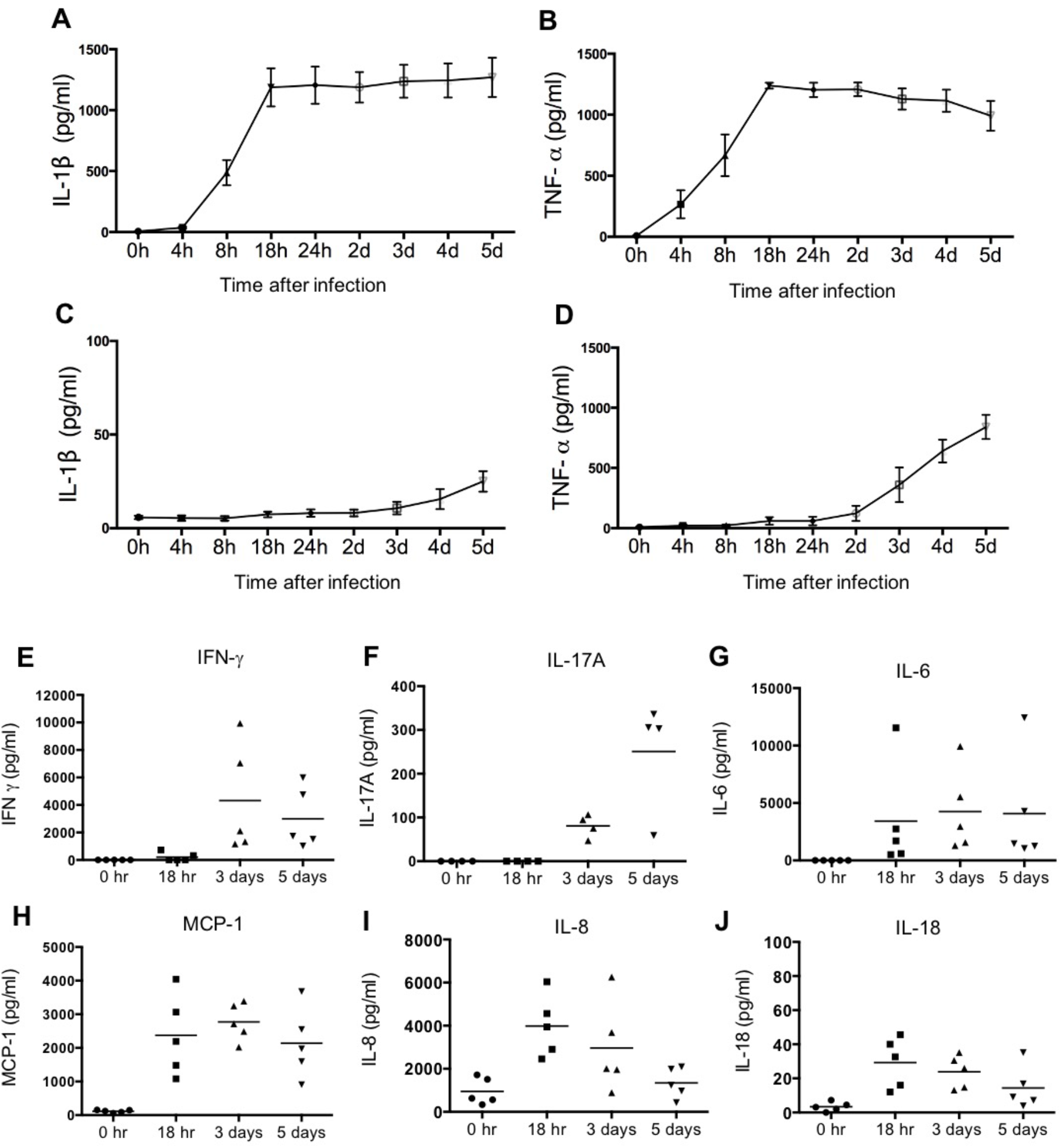
*T. marneffei* yeasts, but not conidia, induce potent IL-1β response in human PBMCs. A, B. Quantification of IL-1β (A) and TNF-α (B) by ELISA in human PBMCs (4×10^6^/ml) infected with live *T. marneffei* yeasts (0.5 MOI) for indicated time points (n=5, mean ± SEM). C, D. ELISA detection of IL-1β (C) and TNF-α (D) in human PBMCs (4×10^6^/ml) infected with live *T. marneffei* conidia (0.5 MOI) for indicated time points (n=5, mean ± SEM). E. Quantification of IFN-γ by LEGENDplex^TM^ in human PBMCs (4×10^6^/ml) infected with live *T. marneffei* yeasts (0.5 MOI) for indicated time points (n=5). F. Quantification of IL-17A by ELISA in human PBMCs (4×10^6^/ml) stimulated with heat-killed *T. marneffei* yeasts (0.5 MOI) for indicated time points (n=4). G-J. Quantitative analyses of cytokines by LEGENDplex^TM^ in human PBMCs (4×10^6^/ml) infected with live *T. marneffei* yeasts (0.5 MOI) for indicated time points (n=5). Data information: In (A-D), “h” and “d” denote “hours” and “days”, respectively. In (E-J), each symbol represents one healthy donor.

To examine whether the viability of *T. marneffei* is a prerequisite for the induction of IL-1β in PBMCs, yeasts and conidia were inactivated by heat (100°C for 40 min) or 4% paraformaldehyde (PFA) treatment for 10 min. Our results showed that there was no significant difference in IL-1β production in PBMCs infected with live, heat-killed or PFA-treated *T. marneffei* yeasts (Fig EV1A), indicating that the viability of *T. marneffei* yeasts was not necessary for the induction of IL-1β response. In contrast, IL-1β production in the PBMCs stimulated with heat-killed conidia was significantly higher compared to live and PFA-treated conidia (Fig EV1B), suggesting that heat treatment might promote the exposure of some pro-inflammatory PAMPs on the conidia cell wall to induce IL-1β response. In addition, both yeast and pseudo-hyphal forms of *C. albicans* were capable of inducing IL-1β production in human PBMCs, and no differential IL-1β response was observed for heat- and PFA-treated *C. albicans* (Fig EV1C and D).

We further evaluated the production of other pro-inflammatory cytokines in *T. marneffei* yeasts-infected human PBMCs. As shown in Fig 1E-J, IFN-γ and IL-17A could be detected only after 3 days of co-culture, whereas IL-6, MCP-1, IL-8 and IL-18 were detectable as early as 18 hr. Specifically, IFN-γ production reached its maximum at 3 days and slightly reduced at 5 days, while IL-17A production greatly increased from 3 days to 5 days of co-culture (Fig 1E and F). IL-6 and MCP-1 production peaked at 18 hr and remained at similar levels at 3 days and 5 days of co-culture (Fig 1G and H). IL-8 and IL-18 production was maximal at 18 hr and showed a downward trend thereafter (Fig 1I and J). These results suggested that *T. marneffei* yeasts sequentially activated innate and adaptive immune responses.

### Differential requirement for IL-1β response to *T. marneffei* yeasts in various human immune cell types

It has been suggested that IL-1β production can be mediated by inflammasome activation which requires two signals: the triggering of pro-IL-1β production (signal 1) and the processing of pro-IL-1β into mature IL-1β (Signal 2) (Latz *et al*, 2013). To investigate whether there was any differential requirement for IL-1β response to yeasts in different cell types, human PBMCs, monocytes, monocytes-derived macrophages and monocytes-derived DCs were primed with or without lipopolysaccharide (LPS) prior to *T. marneffei* yeasts stimulation. As shown in Fig 2A and B, *T. marneffei* yeasts alone were able to induce abundant IL-1β production in PBMCs and monocytes, indicating that yeasts provided ‘signal 1’ and ‘signal 2’ for inflammasome activation in these cell types. In contrast, LPS priming was required for IL-1β response to *T. marneffei* yeasts in macrophages and DCs (Fig 2C and D), suggesting that *T. marneffei* yeasts only provided ‘signal 2’, and the pro-IL-1β induced by LPS (‘signal 1’) was essential in triggering inflammasome activation in these two cell types. On the other hand, high level of TNF-α could be detected in human PBMCs (Fig 2E), monocytes (Fig 2F) and DCs (Fig 2H), but not in macrophages (Fig 2G), after *T. marneffei* yeast stimulation.

**Figure 2.**
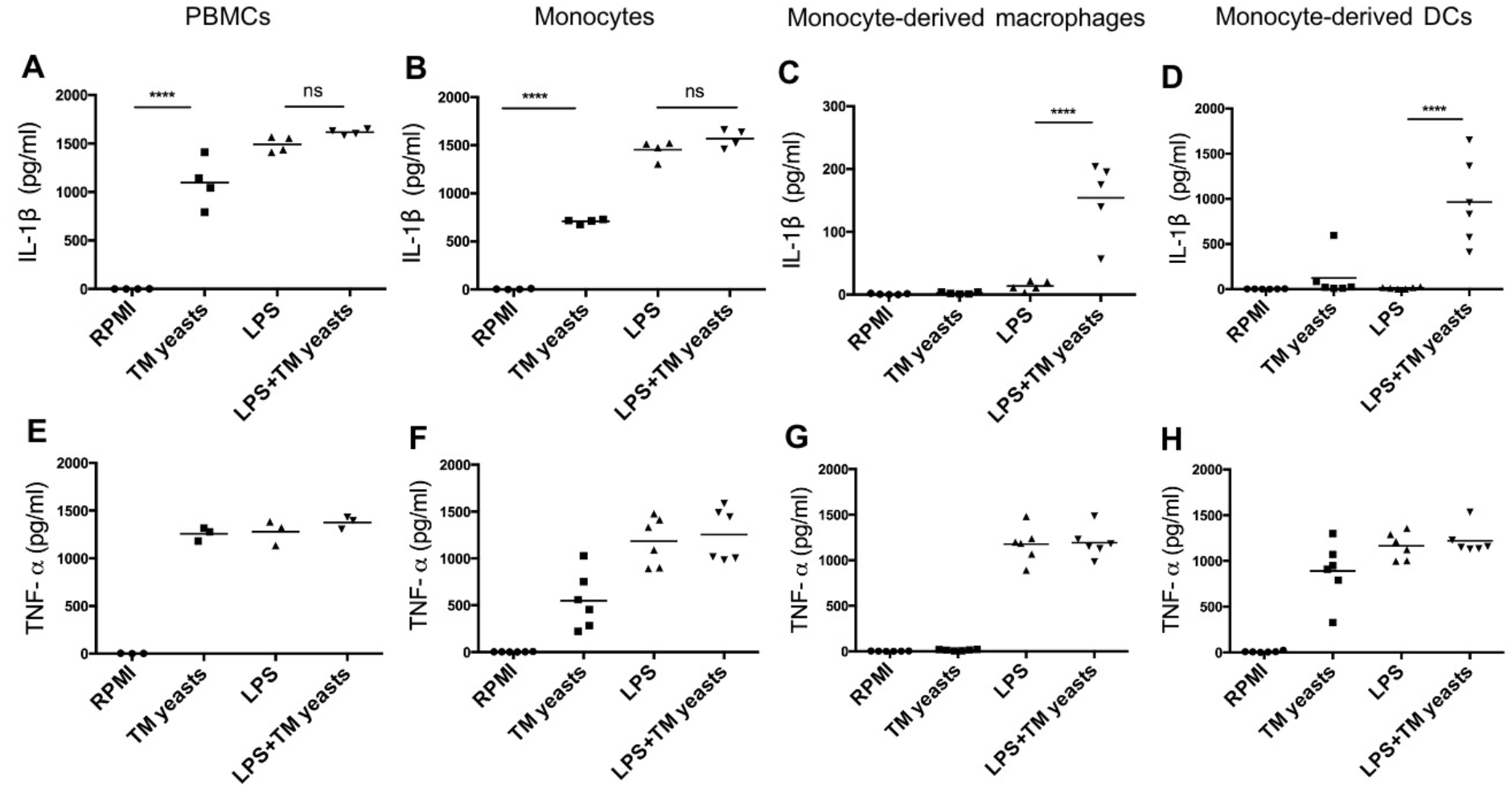
Differential requirement for IL-1β response to *T. marneffei* yeasts in various human immune cell types. A-D. Quantitative ELISA detection of IL-1β in human PBMCs (4×10^6^/ml), CD14^+^ monocytes (2×10^6^/ml), human monocytes-derived macrophages (1×10^6^/ml), CD14^+^ monocytes-derived dendritic cells (DCs) (1×10^6^/ml), respectively, after stimulation with heat-killed *T. marneffei* (TM) yeasts (0.5 MOI) for 18 hr with or without LPS priming. E-H. Quantification of TNF-α by ELISA in human PBMCs (4×10^6^/ml), CD14^+^ monocytes (2×10^6^/ml), human monocytes-derived macrophages (1×10^6^/ml), CD14^+^ monocytes-derived dendritic cells (DCs) (1×10^6^/ml), respectively, after stimulation with heat-killed *T. marneffei* (TM) yeasts (0.5 MOI) for 18 hr with or without LPS priming. Data information: In (A-H), each symbol represents one healthy donor, and data are analyzed by one-way ANOVA. ns =not significant, ****p<0.0001.

### Dectin-1/syk signaling mediates IL-1β response to *T. marneffei* yeasts in human monocytes

Dectin-1 recognizes β-1,3-glucans in fungal cell wall, which leads to pro-inflammatory immune responses (Hardison & Brown, 2012). To elucidate the involvement of Dectin-1 in the initiation of IL-1β response to *T. marneffei* yeasts, human CD14^+^ monocytes were co-cultured with heat-killed *T. marneffei* yeasts in the presence of anti-human Dectin-1 or TLR2 neutralizing antibodies or isotype controls (mouse IgG or human IgA). As shown in Fig 3A, Dectin-1 blockade resulted in a marked reduction of IL-1β in *T. marneffei* yeasts-stimulated monocytes compared with mouse IgG treatment group. TNF-α production was slightly reduced by Dectin-1 blockade, though not significantly different (Fig 3B). These data suggested that Dectin-1 mediated the production of IL-1β and TNF-α in response to *T. marneffei* yeasts in human monocytes. Conversely, blockade of TLR2 resulted in significantly elevated IL-1β secretion and had no effect on TNF-α production in response to yeasts (Fig 3C and D), suggesting that TLR2 negatively regulated IL-1β production in the context of *T. marneffei* stimulation.

**Figure 3.**
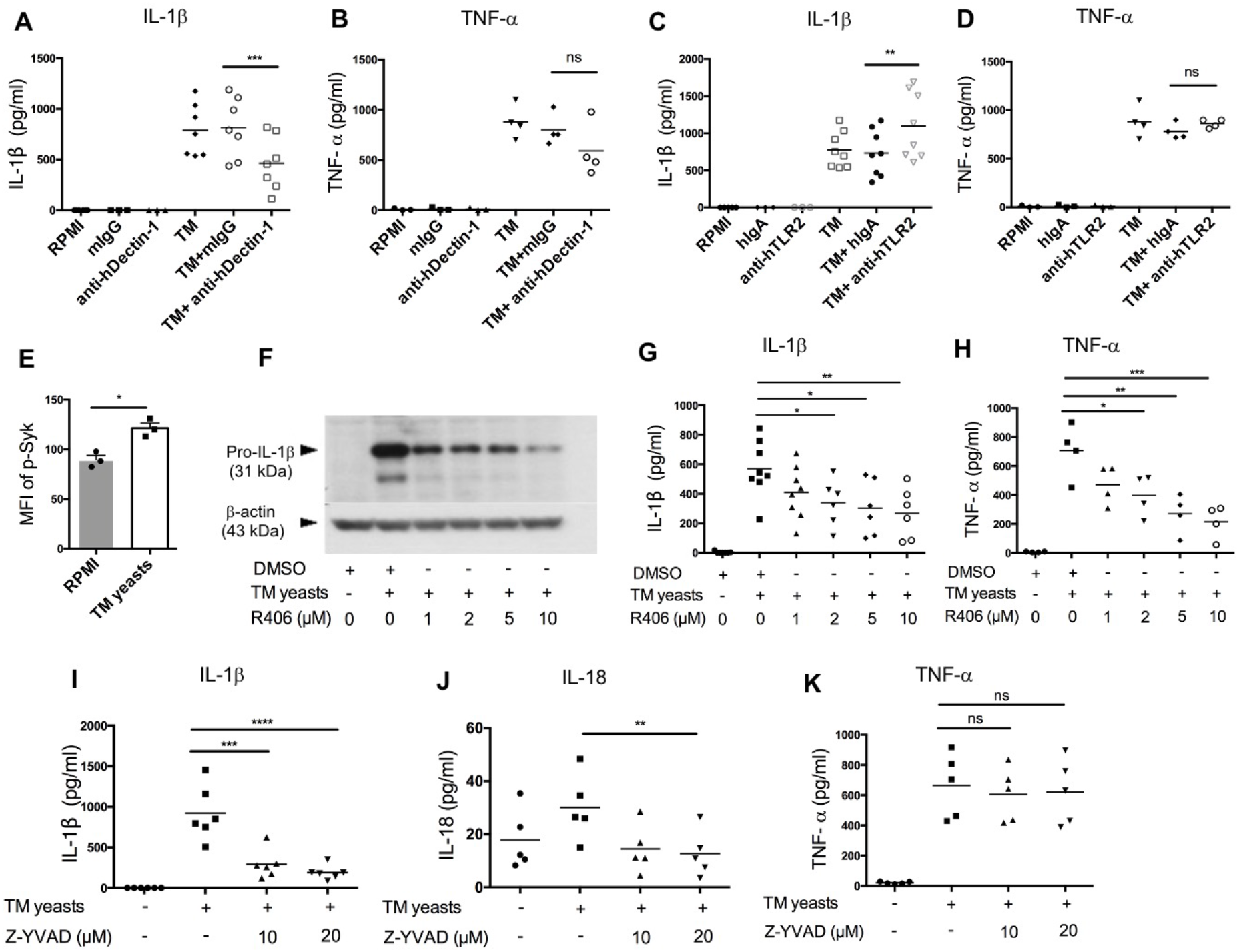
Dectin-1/syk signaling mediates IL-1β response to *T. marneffei* yeasts in human monocytes. A, B. Quantification of IL-1β and TNF-α by ELISA in human CD14^+^ monocytes (2×10^6^/ml) stimulated with heat-killed *T. marneffei* (TM) yeasts (0.5 MOI) for 18 hr in the presence of anti-hDectin-1 blocking antibodies or mouse IgG (mIgG) (10 μg/ml). C, D. Quantification of IL-1β and TNF-α by ELISA in human CD14^+^ monocytes (2×10^6^/ml) stimulated with heat-killed *T. marneffei* (TM) yeasts (0.5 MOI) for 18 hr in the presence of anti-hTLR2 blocking antibodies or human IgA (hIgA) (10 μg/ml). E. Detection of median fluorescence intensity (MFI) of phospho-Syk by flow cytometry in human CD14^+^ monocytes (2×10^6^/ml) stimulated with heat-killed *T. marneffei* (TM) yeasts (0.5 MOI) for 18 hr (n=3, mean ± SEM). F. A representative immunoblot of pro-IL-β of cell lysates in human CD14^+^ monocytes (2×10^6^/ml) stimulated with heat-killed *T. marneffei* (TM) yeasts (0.5 MOI) for 18 hr in the absence or presence of Syk inhibitor, R406 (n=3). G, H. ELISA quantification of IL-1β and TNF-α in the culture supernatant of human CD14^+^ monocytes (2×10^6^/ml) stimulated with heat-killed *T. marneffei* (TM) yeasts (0.5 MOI) for 18 hr in the absence or presence of Syk inhibitor, R406. I-K. Quantification of IL-1β, IL-18 and TNF-α by ELISA in the culture supernatant of human CD14^+^ monocytes (2×10^6^/ml) stimulated with heat-killed *T. marneffei* yeasts (0.5 MOI) for 18 hr in the absence or presence of Z-YVAD (caspase-1 inhibitor). Data information: Data are analyzed by paired two-tailed t test (A-E) or by one-way ANOVA (G-K). Each symbol represents one healthy donor. ns =not significant, *p<0.05, **p<0.01, ***p<0.001, ****p<0.0001.

To study the role of Dectin-1/Syk signaling pathway in the induction of IL-1β response to *T. marneffei* yeasts, we measured Syk phosphorylation in *T. marneffei* stimulated human CD14^+^ monocytes by flow cytometry. As expected, *T. marneffei* yeasts induced Syk phosphorylation compared to the unstimulated cells (Fig 3E). Pre-treatment of CD14^+^ monocytes with R406 (Syk inhibitor) prior to *T. marneffei* yeasts stimulation resulted in significantly reduced expression of pro-IL-1β in cell lysates (Fig. 3F) as well as IL-1β in culture supernatant (Fig 3G), in a dose-dependent manner. A similar trend of dose-dependent inhibition of TNF-α production in monocytes co-cultured with *T. marneffei* yeasts was also observed (Fig 3H). Collectively, our data suggested that Syk, the molecule that acts downstream to Dectin-1 in the initiation of antifungal immunity (Drummond *et al*, 2011), played an important role in eliciting IL-1β response to *T. marneffei* yeasts in human monocytes.

To verify our hypothesis that caspase-1 activation mediates IL-β and IL-18 release in human monocytes infected with *T. marneffei* yeasts, we pre-treated human CD14^+^ monocytes with caspase-1 inhibitor (Z-YVAD, 10μM and 20μM) prior to co-culture with heat-killed *T. marneffei* yeasts. IL-1β and IL-18 secretion was significantly attenuated in Z-YVAD-treated monocytes (Fig 3I and J), whereas inhibition of caspase-1 activity had no obvious effect on TNF-α production (Fig 3K), demonstrating that caspase-1 activation contributed to IL-1β and IL-18 release in response to *T. marneffei*.

### *T. marneffei* yeasts trigger IL-1β production via the NLRP3 inflammasome

To delineate whether IL-1β release induced by *T. marneffei* yeasts was mediated by NLRP3 inflammasome, murine bone marrow-derived DCs (BMDCs) from WT, *Nlrp3*^−/−^, *Casp1*^−/−^ mice were co-cultured with heat-killed yeasts, and then NLRP3, caspase-1 p20 and p45 expression were analyzed by immunoblots. As shown in Fig 4A, we found that NLRP3 expression was obviously upregulated by yeasts in BMDCs from WT and *Casp-1*^−/−^ mice. Moreover, pro-caspase-1 (p45) expression in cell lysates was comparable between unstimulated and yeasts-stimulated cells in either WT or *Nlrp3*^−/−^ mice. Active caspase-1 p20 was detectable in the concentrated supernatant of yeasts-stimulated BMDCs from WT mice, but it was absent in the supernatant from cells deficient in NLRP3 or caspase-1, suggesting that caspase-1 activation is dependent on NLRP3. LPS plus nigericin, which are considered as the activators of classical NLRP3 inflammasome, could activate caspase-1 since active caspase-1 p20 was observed in both cell lysates and supernatant of WT BMDCs.

**Figure 4.**
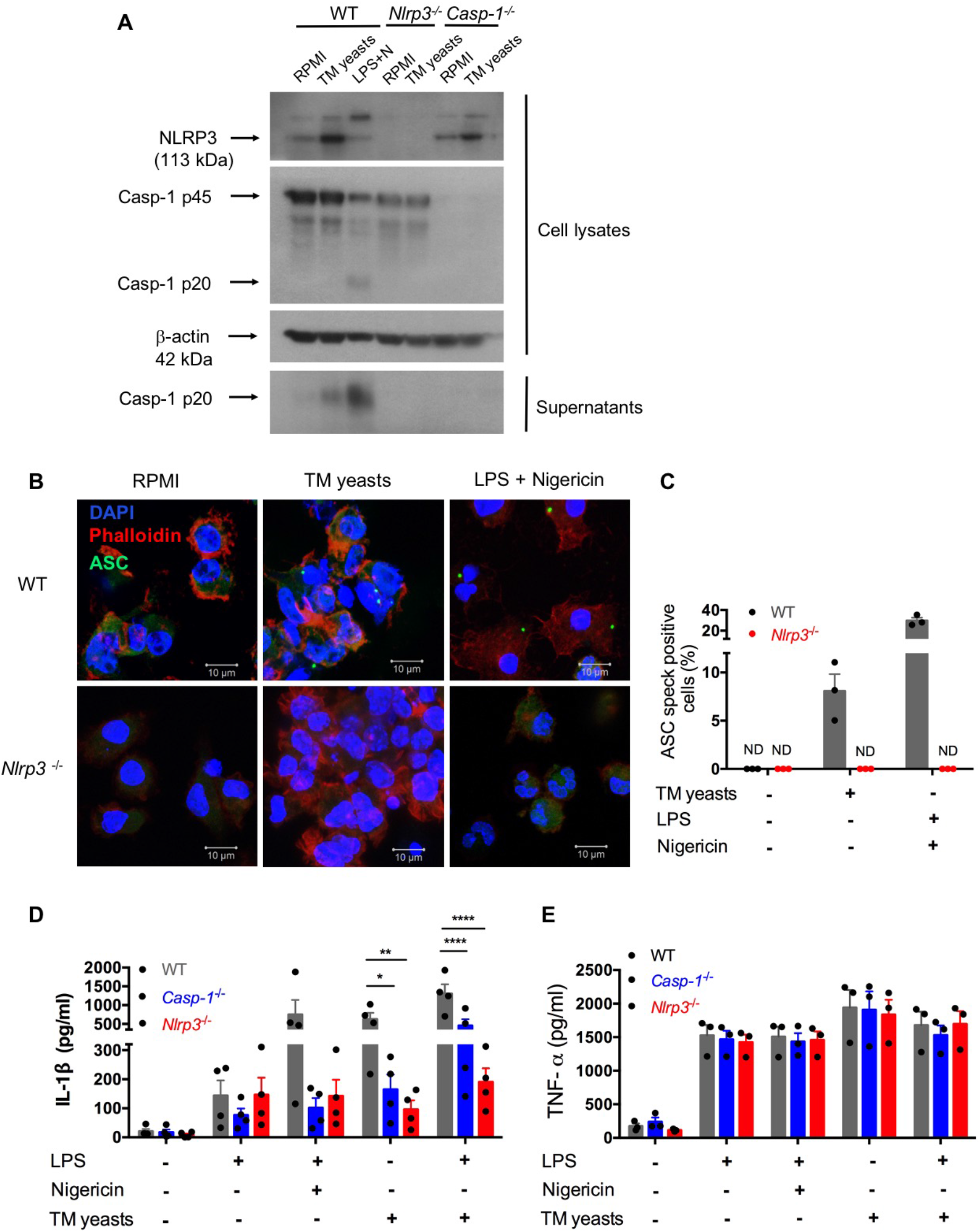
*T. marneffei* yeasts trigger IL-1β production via the NLRP3 inflammasome. A. Representative immunoblots of NLRP3, caspase-1 p45 and p20 from cell lysates and caspase-1 p20 from concentrated cell supernatant in WT, *Nlrp3*^−/−^ and *Casp-1*^−/−^ murine BMDCs stimulated with heat-killed *T. marneffei* (TM) yeasts (1 MOI) for 18 hr, or LPS plus nigericin as a positive control (n=3). B, C. Representative confocal micrographs (B) and the percentages (C) of ASC specks formation in WT, *Nlrp3^−/−^* murine BMDCs (2×10^6^/ml) stimulated with heat-killed *T. marneffei* (TM) yeasts (1 MOI) for 18 hr, or LPS plus nigericin as a positive control. D, E. Quantitative detection of IL-1β and TNF-α by ELISA in the cell supernatant of WT, *Casp-1*^−/−^ and *Nlrp3*^−/−^ murine BMDCs (2×10^6^/ml) stimulated with heat-killed *T. marneffei* (TM) yeasts (1 MOI) for 18 hr with or without LPS priming, or LPS plus nigericin as a positive control. Data information: In (C), the percentages of ASC speck positive cells are quantified and depicted as mean ± SEM of n=3 mice for each group, and “ND” denotes “not detectable”. In (D, E), data represent mean±SEM of n=4 mice, and data are analyzed by two-way ANOVA. *p<0.05, **p<0.01, ****p<0.0001.

To determine whether *T. marneffei* yeasts could induce ASC specks to mediate IL-1β processing, BMDCs from WT and *Nlrp3*^−/−^ mice were co-cultured with yeasts, and the formation of ASC pyroptosomes (“specks”) was examined. Confocal microscopy revealed that ASC pyroptosomes, appearing as a single cytoplasmic speck, was present in WT cells but not in *Nlrp3*^−/−^ cells stimulated with *T. marneffei* yeasts or LPS plus nigericin (Fig 4B). We also quantified the percentage of cells containing ASC specks and observed that around 8% of WT cells formed ASC specks while it was undetectable in *Nlrp3*^−/−^ cells after *T. marneffei* yeasts stimulation (Fig 4C). Similarly, LPS plus nigericin could induce about 29% of WT cells containing ASC specks (Fig 4C). Our results indicated that *T. marneffei* yeasts induced ASC speck formation, which was dependent on NLRP3, to induce IL-1β processing.

Next, to further test the hypothesis that IL-1β response to *T. marneffei* yeasts was dependent on NLRP3 inflammasome, BMDCs from WT, *Casp-1^−/−^*, and *Nlrp3*^−/−^ mice were co-cultured with heat-killed *T. marneffei* yeasts, and IL-1β and TNF-α in the supernatant were examined. As shown in Fig 4D, LPS induced a small amount of IL-1β in these BMDCs, whereas there was remarkably lower IL-1β in *Casp-1^−/−^* and *Nlrp3*^−/−^ BMDCs than WT cells after stimulation with LPS plus nigericin. More importantly, we observed that BMDCs from *Casp-1^−/−^* and *Nlrp3*^−/−^ mice had significantly impaired IL-1β response to *T. marneffei* yeasts when compared with WT mice (Fig 4D). Furthermore, an intriguing observation was that IL-1β production in BMDCs from *Nlrp3*^−/−^ mice was slightly lower than those from *Casp-1^−/−^*mice after *T. marneffei* yeasts stimulation with or without LPS priming (Fig 4D), indicating that IL-1β response to *T. marneffei* yeasts was more dependent on NLRP3 than caspase-1. In contrast to IL-1β production, BMDCs from WT, *Casp-1*^−/−^, and *Nlrp3*^−/−^ mice produced comparable levels of TNF-α when co-cultured with *T. marneffei* yeasts (Fig 4E). Collectively, these results corroborated that NLRP3 and caspase-1 were required for IL-1β response to *T. marneffei* yeasts in murine BMDCs.

### *T. marneffei* yeasts elicit differential Th1 and Th17 immune responses in human CD14^+^ monocytes and monocytes-derived DCs

We have observed that *T. marneffei* yeasts triggered IFN-γ and IL-17A production in human PBMCs (Fig 1E and F). To further validate our hypothesis that *T. marneffei* yeasts would induce Th1 and Th17 immune responses, isolated human CD4^+^ T cells were co-cultured with autologous monocytes or monocytes-derived DCs with *T. marneffei* yeasts or *C. albicans* for 7 and 12 days, and intracellular IFN-γ and IL-17A production in CD4^+^ T cells was quantified by flow cytometry. Upon co-culture of CD4^+^ T cells with *T. marneffei* yeasts, no IFN-γ- or IL-17A-producing cells could be detected (Figs 5A-C, and EV2A and B). However, when CD4^+^ T cells were co-cultured with *T. marneffei* yeasts for 7 and 12 days in the presence of monocytes, robust IFN-γ and/or IL-17A production was evident (Figs 5A,B, G, and EV2C and D), while no statistically significant differences were observed for IL-17A-producing cells on day 12 (Fig 5B) and IFN-γ^+^IL-17A^+^-producing cells on day 7 and 12 (Fig 5C) after stimulation with *T. marneffei* yeasts. These findings suggested that the induction of Th1 and Th17 response by *T. marneffei* yeasts required the presence of antigen presenting cells (APCs) such as monocytes. We also found that *C. albicans* yeasts induced much stronger Th1 and Th17 immune responses than *T. marneffei* in the presence of monocytes (Figs 5A-C, and G, and EV2C and D). Consistent with IFN-γ and IL-17A production, increased T-bet and RORγt, the master transcription factors of respective Th1 and Th17 cells, were observed in CD4^+^ T cells after co-culture with TM yeasts-primed autologous monocytes for 7 days (Fig EV2G and H). Distinct from CD14^+^ monocytes, monocytes-derived DCs stimulated with *T. marneffei* or *C. albicans* yeasts induced delayed Th1 and Th17 cell differentiation; the remarkable increase in the percentage of IFN-γ -and/or IL-17A-producing CD4^+^ T cells could mostly be detected on day 12 when T cells were co-cultured with *T. marneffei* or *C. albicans* yeasts in the presence of DCs (Figs 5D-F, and EV2E and F).

**Figure 5.**
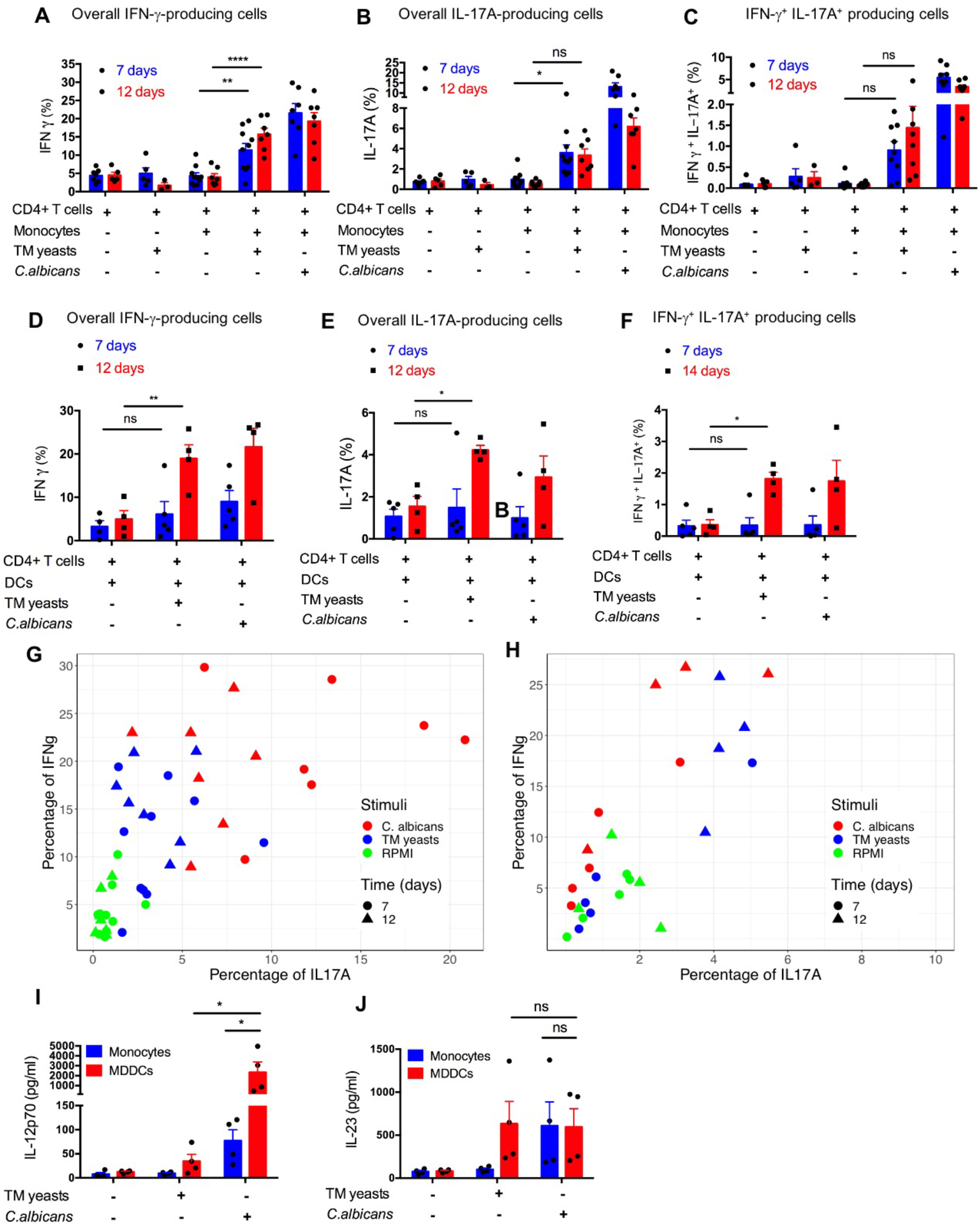
*T. marneffei* yeasts elicit differential Th1 and Th17 immune responses in human CD14^+^ monocytes and monocytes-derived DCs. A-C. Analyses of IFN-γ and/or IL-17A producing CD4 T cells by flow cytometry in human CD4^+^ T cells (1×10^5^/well) co-cultured with heat-killed *T. marneffei*- or *C. albicans*-stimulated autologous CD14^+^ monocytes (1×10^5^/well) for 7 days and 12 days. D-F. Analyses of IFN-γ and/or IL-17A producing CD4 T cells by flow cytometry in human CD4^+^ T cells (1×10^5^/well) co-cultured with heat-killed *T. marneffei*- or *C. albicans*-stimulated autologous monocytes-derived DCs (0.2×10^5^/well) for 7 days and 12 days. G. Scatter plot of overall IFN-γ-producing CD4 T cells (y axis) and overall IL-17A-producing CD4 T cells (x axis) in (A and B). H. Scatter plot of overall IFN-γ-producing CD4 T cells (y axis) and overall IL-17A-producing CD4 T cells (x axis) in (D and E). I. Quantitative ELISA analysis of IL-12p70 in the supernatant of human CD14^+^ monocytes (Monocytes) (2×10^6^/ml) and monocytes-derived DCs (MDDCs) (1×10^6^/ml) stimulated with *T. marneffei* yeasts or *C. albican* yeasts for 18 hr. J. Quantitative ELISA analysis of IL-23 in the supernatant of human CD14^+^ monocytes (Monocytes) (2×10^6^/ml) and monocytes-derived DCs (MDDCs) (1×10^6^/ml) stimulated with *T. marneffei* yeasts or *C. albican* yeasts for 18 hr. Data information: In (A-H), data are given as the percentage of total gated CD3^+^ cells. In (A-F, and I-J), data are analyzed by two-way ANOVA and depicted as means ± SEM. ns =not significant, *p<0.05, **p<0.01, p<0.0001. Each symbol represents one healthy donor in all the data.

In order to further dissect the difference between monocytes and DCs in promoting the differentiation of CD4^+^ T cells after fungal stimulation, we next assessed the release of IL-12p70 and IL-23, which are considered to be the key cytokines driving Th1 and Th17 cell differentiation respectively, from human CD14^+^ monocytes and monocyte-derived DCs. We found that the induction of IL-12p70 and IL-23 by *T. marneffei* was negligible in monocytes (Fig 5I and J). In contrast, IL-23 was induced by *T. marneffei* in DCs at a level comparable with *C. albicans* (Fig 5J).

Taken together, our data showed that *T. marneffei* yeasts elicited Th1 and Th17 immune responses in the presence of APCs. CD14^+^ monocytes exhibited earlier induction of IFN-γ-producing and IL-17A-producing CD4^+^ T cells than monocyte-derived DCs when co-cultured with *T. marneffei* or *C. albicans*.

### Caspase-1 inhibition attenuates Th1 and Th17 immune response to *T. marneffei* yeasts

To characterize the role of activated NLRP3 inflammasome in the Th1 and Th17 immune responses towards *T. marneffei* yeasts, human PBMCs were co-cultured with *T. marneffei* yeasts for 5 days in the presence of caspase-1 inhibitor, Z-YVAD. Blocking of caspase-1 activation resulted in significantly reduced IFN-γ and IL-17A release in response to *T. marneffei* yeasts (Fig 6A). Next, we determined the percentage of IFN-γ- and/or IL-17A-producing CD4^+^ T cells upon co-culture of *T. marneffei* yeasts-stimulated monocytes and CD4^+^ T cells in the presence of Z-YVAD. Our results showed that caspase-1 inhibition led to approximately 2-fold reduction in overall IL-17A-producing CD4^+^ T cells (Fig 6B and D) and IFN-γ/IL-17A-producing CD4^+^ T cells (Fig 6B and E), although there was no significant effect on the overall intracellular IFN-γ production (Fig 6B and C). Our data indicated that NLRP3 inflammasome activation partially enhanced *T. marneffei* yeasts-induced Th1 and Th17 immune responses in human immune cells *in vitro*.

**Figure 6.**
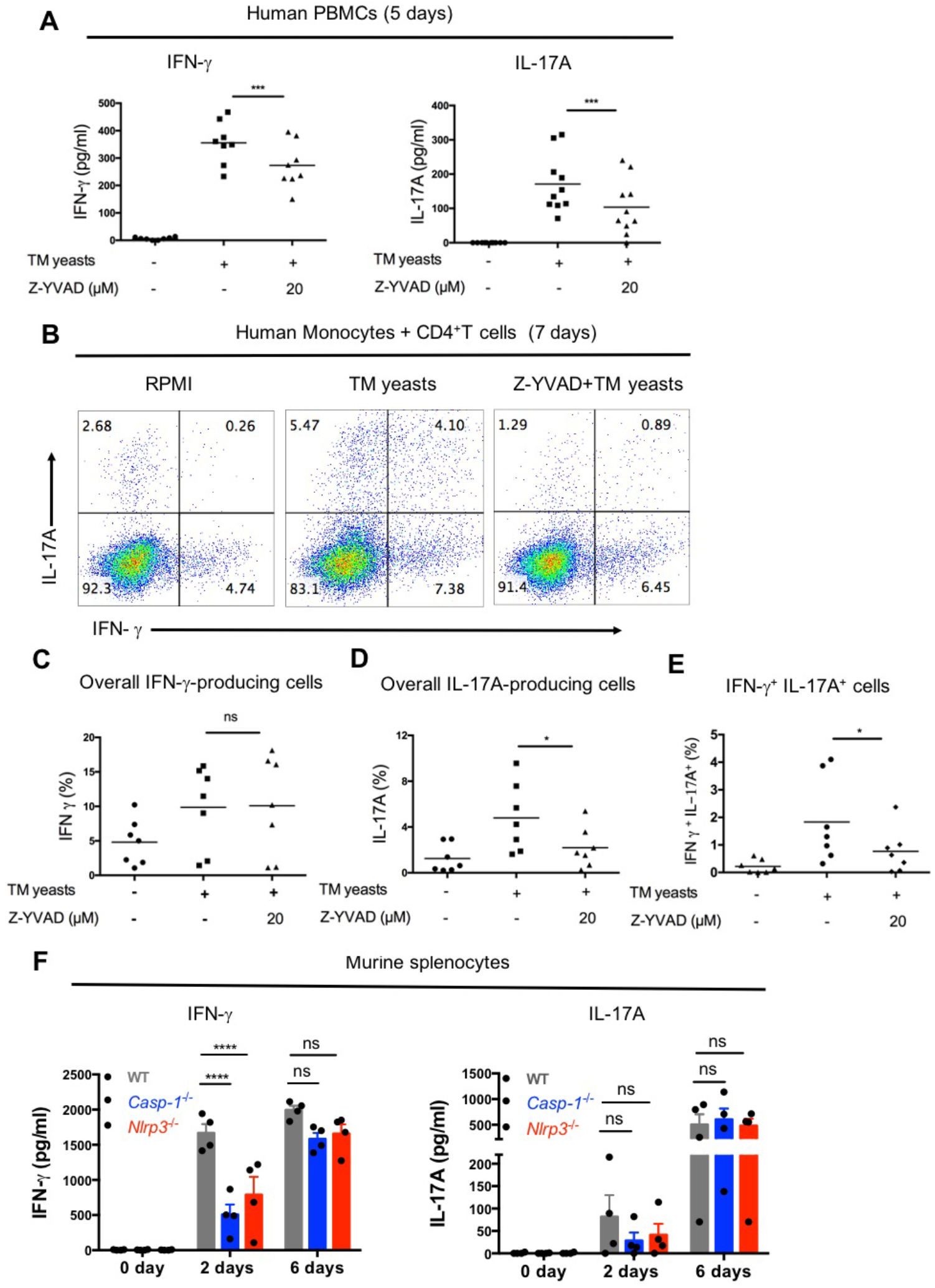
Caspase-1 inhibition attenuates Th1 and Th17 immune responses to *T. marneffei* yeasts. A. Quantification of IFN-γ and IL-17A by ELISA in the supernatant of human PBMCs (4×10^6^/ml) stimulated with heat-killed *T. marneffei* (TM) yeasts (0.5 MOI) in the absence or presence of caspase-1 inhibitor, Z-YVAD (20μM), for 5 days. B. Representative plots of intracellular IFN-γ and/or IL-17A in the CD4^+^ T cells (1×10^5^/well) co-cultured with heat-killed *T. marneffei* (TM) yeasts-stimulated autologous CD14^+^ monocytes (1×10^5^/well) in the absence or presence of Z-YVAD (20μM) for 7 days. C-E. Statistical analyses of overall IFN-γ-producing cells (C) and overall IL-17A-producing cells (D) and IFN-γ^+^IL-17A^+^-producing cells (E) in co-cultured CD4^+^ T cells and *T. marneffei*-primed autologous CD14^+^ monocytes in the absence or presence of Z-YVAD for 7 days as described in (B). F. Detection of murine IFN-γ and IL-17A by ELISA in the supernatant of WT, *Casp-1^−/−^* and *Nlrp3^−/−^* murine splenocytes (2×10^6^/ml) stimulated with heat-killed *T. marneffei* yeasts (1 MOI) for indicated time points. Data information: In (A) and (C-E), each symbol represents one healthy donor, and data are analyzed by paired two-tailed Student’s t tests. In (F), data are depicted as mean ± SEM of n=4 mice and are analyzed by two-way ANOVA. ns=not significant, *p<0.05, ***p<0.001, ****p<0.0001.

To clarify the role of caspase-1 and NLRP3 in the induction of Th1 and Th17 response to *T. marneffei*, we isolated murine splenocytes from wild-type, *Casp-1*^−/−^, *Nlrp3*^−/−^ mice and incubated them with *T. marneffei* yeasts for 2 days or 6 days, and the supernatant was collected for the quantification of IFN-γ and IL-17A. As shown in Fig 6F, it was observed that splenocytes from *Casp-1*^−/−^ and *Nlrp3*^−/−^ mice had impaired IFN-γ and IL-17A production compared to WT mice at 2 days post stimulation, while there was no statistically significant effect on IL-17A. On day 6 of incubation, IFN-γ and IL-17A production by *Casp-1*^−/−^ and *Nlrp3*^−/−^ splenocytes was sharply increased and reached a similar level as splenocytes from WT mice. Our findings suggested that NLRP3 inflammasome activation promoted early stage of IFN-γ production in murine splenocytes.

### The NLRP3 inflammasome controls antifungal immunity *in vivo*

Our *in vitro* study has demonstrated that *T. marneffei* yeasts activated NLRP3 inflammasome that promoted Th1 and Th17 immune responses. To examine the protective role of NLRP3 inflammasome in defense against *T. marneffei* infection *in vivo*, WT, *Casp-1*^−/−^, *Nlrp3*^−/−^ mice were infected with 5×10^5^ CFU of *T. marneffei* yeasts per mouse by intravenous injection and their survival was monitored. As shown in Fig 7A, *Casp-1*^−/−^ and *Nlrp3*^−/−^ mice were more susceptible to *T. marneffei* yeasts infection than WT mice. Of note, *Casp-1*^−/−^ mice showed more resistance to fungal infection than *Nlrp3*^−/−^ mice. These observations suggested that NLRP3 and caspase-1 conferred host protection against *T. marneffei* infection, in particular, NLRP3 played a more protective role than caspase-1. To further investigate the factors resulting in the differential survival rates among WT, *Casp-1*^−/−^, *Nlrp3*^−/−^ mice, murine spleens and livers were collected at 7 and 14 days post infection (dpi) for fungal load detection. There was comparable fungal load in spleens from WT, *Casp-1*^−/−^, *Nlrp3*^−/−^ mice at 7 dpi (Fig 7B). Nevertheless, significantly elevated fungal load was observed in spleens from *Nlrp3*^−/−^ mice when compared to WT mice at 14 dpi (Fig 7B). Similarly, liver specimen from *Nlrp3*^−/−^ mice exhibited the highest fungal load, followed by *Casp-1*^−/−^ mice, and WT mice showed the least fungal load in livers at 7 and 14 dpi (Fig 7C). It was worth noting that the fungal load in spleens and livers from WT mice at 14 dpi was similar to that at 7 dpi, whereas the fungal load in these organs from *Nlrp3*^−/−^ mice at 14 dpi was significantly higher than that 7 dpi (Fig 7B and C), indicating that NLRP3 inflammasome played a pivotal role in the control of *T. marneffei* proliferation.

**Figure 7.**
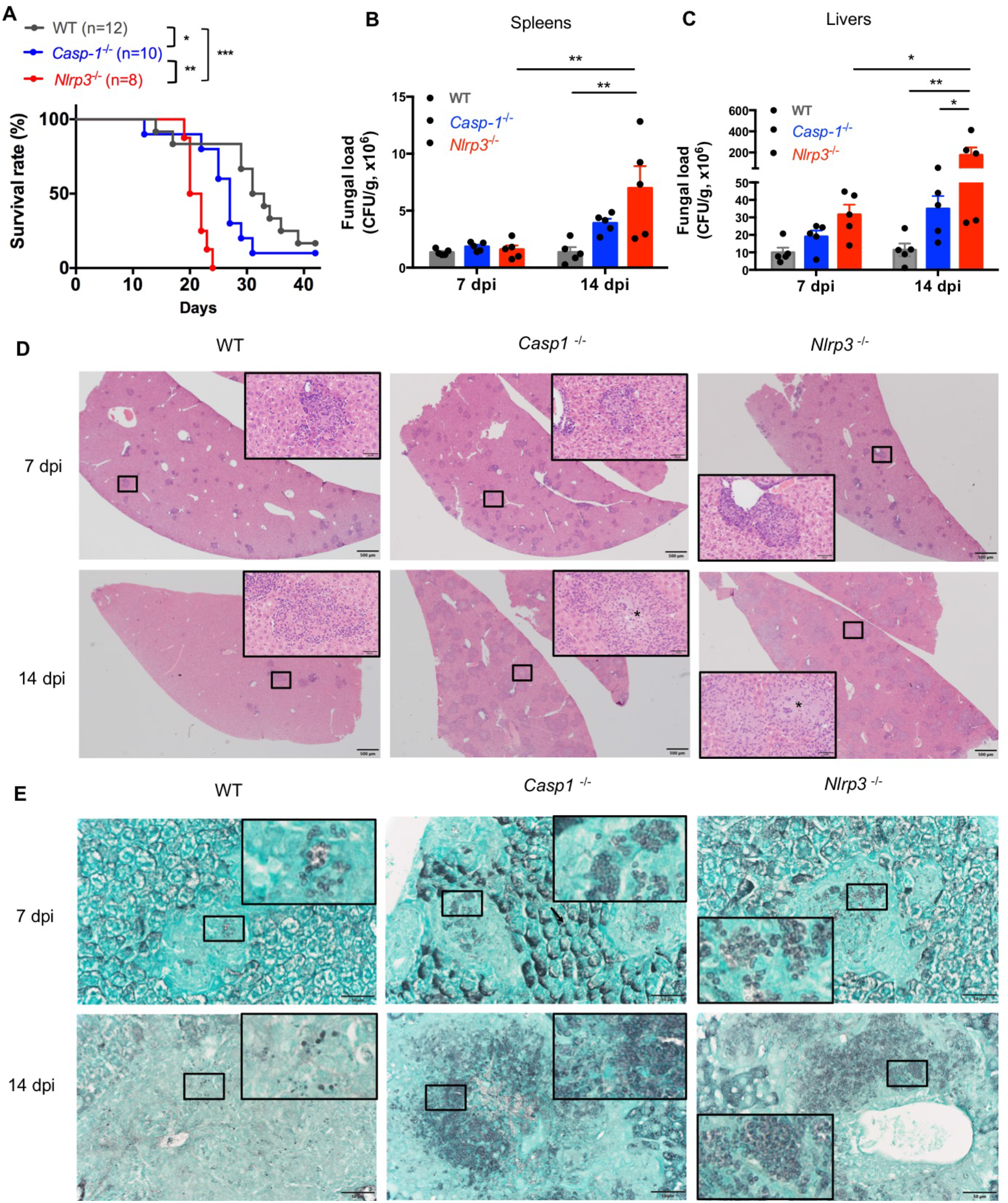
The NLRP3 inflammasome controls antifungal immunity in vivo. A. Survival plot of WT, *Casp-1*^−/−^ and *Nlrp3*^−/−^ mice intravenously injected with live *T. marneffei* yeasts (5×10^5^ CFU per mouse). B, C. Fungal load analyses of spleens (B) and livers (C) of WT, *Casp-1*^−/−^ and *Nlrp3*^−/−^ mice infected with live *T. marneffei* yeasts at 7 and 14 days post infection (dpi). D. Representative HE staining graphs of murine livers of WT, *Casp-1*^−/−^ and *Nlrp3*^−/−^ mice infected with live *T. marneffei* yeasts (n=4). Scale bar denotes 500 μm. The insets indicate the magnified area of the smaller box containing granulomas, and the asterisk indicates massive fungal yeasts in the granulomas. E. Representative GMS staining graphs of murine livers of WT, *Casp-1*^−/−^ and *Nlrp3*^−/−^ mice infected with live *T. marneffei* yeasts (n=4). Scale bar denotes 50 μm. The insets indicate the magnified area of the smaller box containing *T. marneffei* yeasts. Data information: In (A), the log-rank tests are performed when compared between WT and *Nlrp3*^−/−^ mice groups, and between *Nlrp3*^−/−^ and *Casp-1*^−/−^ groups; the Gehan-Breslow-Wilcoxon test is performed between WT and *Casp-1*^−/−^ groups. In (B and C), data are depicted as mean ± SEM of n=5 mice and are analyzed by two-way ANOVA. *p<0.05, **p<0.01, ***p<0.001.

Histological studies of liver sections from WT, *Casp-1*^−/−^ and *Nlrp3*^−/−^ mice at 7 dpi showed a great number of granulomas (Fig 7D, upper panel). By day 14, granulomas were considerably enlarged and almost replaced the liver parenchyma in *Casp-1*^−/−^ and *Nlrp3*^−/−^ mice when compared to those at 7 dpi, but liver sections from WT mice exhibited a smaller number of granulomas (Fig 7D, lower panel). HE staining of livers revealed that WT mice had relatively fewer cytoplasmic fungal yeasts within macrophages in the central granulomas than *Casp-1*^−/−^ and *Nlrp3*^−/−^ mice (Fig 7D, insets of lower panel). Tissue necrosis was not observed in all three strains of mice. GMS staining further verified that fungal yeasts were present in the center of granulomas (Fig 7E). Slightly fewer yeasts were observed in WT mice than *Nlrp3*^−/−^ and *Casp-1*^−/−^ mice at 7 dpi (Fig 7E, upper panel). Importantly, massive number of *T. marneffei* yeasts were seen in the granulomas of *Casp-1*^−/−^ and *Nlrp3* ^−/−^ livers, while *T. marneffei* yeasts were sparse in the granulomas of WT mice at 14 dpi (Fig 7E, lower panel). Spleen sections of *Nlrp3*^−/−^ mice showed a more severe loss of follicular structure in the white pulp than WT and *Casp*-1^−/−^ mice at 7 dpi (Fig EV3A, upper panel). No granulomas were observed in the spleens (Fig EV 3A). We failed to detect *T. marneffei* in the spleens at 7 dpi, but scattered fungal yeasts could be seen in the spleens of WT, *Nlrp3*^−/−^ and *Casp-1*^−/−^ mice at 14 dpi by GMS staining (Fig EV3B). In conclusion, these histopathological examinations further demonstrated that NLRP3 inflammasome was essential for host defense against *T. marneffei* infection *in vivo*.

## Discussion

*T. marneffei* is an important form of endemic mycoses causing major morbidity and mortality in patients with HIV infection, as well as those with primary and secondary immunodeficiencies. However, innate immune recognition of *T. marneffei* and the induction of adaptive immunity, is largely unknown. For the first time, our study characterizes Syk- and caspase-1 mediated IL-1β response towards *T. marneffei* in human myeloid cells, and the role of caspase-1 in the induction of Th1 and Th17 response in human lymphocytes. Importantly, our murine model of disseminated talaromycosis demonstrates increased fungal load in the liver and spleen of *Casp-1*^−/−^ and *Nlrp3*^−/−^ mice, and correlates with reduced survival.

Human *T. marneffei* infection is initiated by inhalation of conidia existing in the environment. After phagocytosis by pulmonary alveolar macrophages, *T. marne*ffei conidia undergo morphological switch and germinate into the pathogenic yeast cells, which readily disseminate throughout the body causing systemic infection. Our findings reveal differential cytokine response towards *T. marneffei* conidia and yeasts in human PBMCs (Fig 1A-D). These interesting findings indicate that conidia are less immunological than yeasts, as shown by negligible IL-1β and TNF-α production in PBMCs during the first 24-48 hours of co-culture. The delayed rise in TNF-α, and IL-1β to a lesser extent, after 3 days of co-culture probably represents the switch of conidia to yeast form *in vitro*, suggesting that the transformation of conidia into yeasts would be required for IL-1β response. The change in cell wall composition and architecture during different life stages of fungi can lead to differential exposure of PAMPs in the inner cell wall, causing varying degree of PRR activation which determines the level of pathogenicity (Cheng *et al*, 2011).

Our work demonstrates that IL-1β response to *T. marneffei* yeasts is partially mediated by Dectin-1/Syk signaling pathway. Blockade of Dectin-1 with neutralizing antibodies impairs the production of IL-1β and TNF-α to a lesser extent, in response to *T. marneffei* yeasts in human CD14^+^ monocytes. This suggests that Dectin-1 plays a role in initiating downstream signaling that leads to pro-IL-1β and TNF-α production, which has been demonstrated in other fungal pathogens such as *C. albicans* and *H. capsulatum* (Chang *et al*, 2017; Hise *et al*, 2009). We further show that *T. marneffei* yeasts are capable of inducing Syk kinase phosphorylation, and blockade of Syk kinase significantly inhibits both pro-IL-1β and IL-1β as well as TNF-α in a dose-dependent fashion (Fig 3F-H). Acquired immunity to other dimorphic fungi such as *Blastomyces dermatitidis, Histoplasma capsulatum* and *Coccidioides posadasii* infection variably depends on innate sensing by Dectin-1, Dectin-2 and Mincle (Wang *et al*, 2014), and the role of these CTL receptors in cytokine response against *T. marneffei* will require further studies.

TLR2 heterodimerizes with either TLR1 or TLR6 and recognizes triacylated and diacylated lipoprotein. TLR2 and dectin-1 physically associate with one another and synergize to augment anti-fungal response by modulating cytokine production (Dennehy *et al*, 2009; Gantner *et al*, 2003; Hise *et al*, 2009). Our results show that blockade of TLR2 leads to increased IL-1β production but has no impact on TNF-α production in *T. marneffei* yeasts-stimulated human monocytes (Fig 3C and D), suggesting that TLR2 ligation might exert a negative effect on IL-1β-mediated immune response against *T. marneffei.* TLR2-deficient mice infected with *P. brasiliensis* have preferential activation of Th17 response and lower fungal load, while Treg expansion is diminished and aggravates lung inflammation (Loures *et al*, 2010). A similar role of TLR2 was also observed in a murine model of disseminated candidiasis, where TLR2 exerts anti-inflammatory effect by promoting IL-10 production and Treg cell proliferation (Netea *et al*, 2004).

It was previously shown that IL-12p40 production is induced by *T. marneffei* in murine BMDCs, and is mediated by Dectin-1, TLR2 and MyD88 (Nakamura *et al*, 2008). Our data showed that IL-12p70 could not be induced by *T. marneffei* yeasts in human CD14^+^ monocytes and monocyte-derived DCs. This confirms the previous published finding that *T. marneffei* yeasts do not upregulate IL-12 expression in human monocyte-derived DCs and monocyte-derived macrophages (Ngaosuwankul *et al*, 2008). Furthermore, our findings provide strong evidence that human Th17 cells are expanded when co-cultured with *T. marneffei*-stimulated monocytes or monocyte-derived DCs, in contrast to the findings from a recent study showing that murine BMDCs are unable to induce, or actually lower, the percentage of Th17 cells upon co-culture with *T. marneffei* (Tang *et al*, 2020). These observations point to the possible difference in Th1 and Th17 immune response towards *T. marneffei* in human vs murine hosts.

We demonstrate that IFN-γ and IL-17A-producing human CD4^+^ T cells are induced by *T. marneffei*-primed human CD14^+^ monocytes (Fig 5A and B and G), albeit less strong as *C. albicans*-primed monocytes. IL-12 is an important cytokine which induces the differentiation of naïve CD4^+^ T cell into IFN-γ secreting Th1 cells, whereas differentiation of naïve CD4^+^ T-cells to Th17 cells is a result of the combined activity of IL-23 and IL-1β (Goncalves *et al*, 2017; Zhou *et al*, 2009). As neither IL-12 nor IL-23 could be detectable upon co-culture of *T. marneffei*-stimulated human CD14^+^ monocytes (Fig 5I and J), this suggests that cytokines other than IL-12 and IL-23 might be secreted by human monocytes for the differentiation of Th1 and Th17 cells in response to *T. marneffei* infection. For example, it was found that IL-6 contributes to IFN-γ expression in lung tissues in a murine model of systemic *P. brasiliensis* infection (Tristão *et al*, 2017). Alternatively, Th17 cells against *T. marneffei* could originate from memory T cells that are cross-reactive to *C. albicans* instead of being differentiated *de novo*, since Th17 cells cross-reactive to *C. albicans* have been demonstrated to contribute to Th17 response towards *A. fumigatus* (Bacher *et al*, 2019).

On the other hand, IL-23 could be induced by *T. marneffei* yeasts in monocyte-derived DCs, but not IL-12 (Fig 5J). The preferential induction of IL-23 over IL-12 response in DCs is possibly related to dectin-1-mediated signaling, as previously shown in *H. capsulatum* (Chamilos *et al*, 2010). Furthermore, IL-23 is induced by *T. marneffei* in human DCs at a level comparable to *C. albicans*, and correlates with a similar percentage of both IFN-γ and IL-17A-producing CD4^+^ T-cells upon co-culture with the two fungal pathogens. IL-12 and IL-23 are thought to promote mutually exclusive CD4^+^ Th1 and Th17 cell fates (Teng *et al,* 2015), However, recent investigations on human monogenic IL12Rβ2 and IL-23R deficiency, which result in defective responses to IL-12 or IL-23 respectively, show that the isolated absence of IL-12 or IL-23 could in part be compensated by its counterpart for the production of IFN-γ, conferring protection against intracellular pathogen such as mycobacteria (Martínez-Barricate *et al*, 2018). In the absence of IL-12, IL-17 is the requisite for the protective effect induced by IL-23 in pulmonary histoplasmosis (Deepe Jr. & Gibbons, 2009). This might explain how loss-of-function mutation in *STAT3*, which encodes the transcription factor activated by IL-23, leads to increased susceptibility to dimorphic fungi such as *T. marneffei*, *C. immitis* and *H. capsulatum* in AD hyper-IgE syndrome (Lee & Lau, 2017).

IL-1β is essential for Th17 differentiation in human and mice (Acosta-Rodriguez *et al*, 2007; Chung *et al*, 2009). IL-1β synergizes with IL-23 in controlling the expression of RORγt and IL-23 receptor to sustain IL-17 production by effector Th17 cells (Chung *et al*, 2009). Another report also indicates that NLRP3 inflammasome-derived IL-18 and IL-1β drive protective Th1 and Th17 immune responses in disseminated candidiasis (van de Veerdonk *et al*, 2011). Consistently, treatment of human PBMCs with caspase-1 inhibitor, which suppressed the production of IL-1β and IL-18 production in human monocytes (Fig 3I and J), significantly impaired the secretion of IFN-γ and IL-17A, and the differentiation of IL-17A-producing-T cells (Fig 6C-E). These findings highlight the important role of caspase-1 in bridging the innate and adaptive immune response, particularly Th17 response towards *T. marneffei* infection in human cells.

Furthermore, in line with the role of inflammasome activation in host defense against *C. albicans* (Hise *et al,* 2009) and *A. fumigatus* (Karki *et al,* 2015), our study defined the critical role of NLRP3 and caspase-1 in the induction of IL-1β response to *T. marneffei*. In WT BMDCs, *T. marneffei* yeasts strongly upregulated the expression of NLRP3, cleavage of pro-caspase-1 to caspase-1 p20, and the formation of ASC pyroptosomes (Fig 4A and C). Using the murine model of systemic *T. marneffei* infection, we found that *Nlrp3*^−/−^ and *Casp-1*^−/−^ mice had higher mortality rate. Of note, the fungal load in the spleen and liver of WT mice were comparable on 7 dpi and 14 dpi, while an increase in fungal load became obvious on 14 dpi in *Casp-1*^−/−^ and *Nlrp3*^−/−^ mice, particularly the latter (Fig 7A-C). This suggests that NLRP3/Caspase-1 signaling is crucial in controlling fungal proliferation in vivo.

An intriguing observation is that the significantly lower survival rate of *Nlrp3*^−/−^ mice correlated with higher fungal load, particularly in the liver at 14 dpi, compared with *Casp-1*^−/−^ mice (Fig 7A-C). These *in vivo* data correlate well with IL-1β response to *T. marneffei* in murine BMDCs *in vitro*, reinforcing the critical role of IL-1β in protective immunity against *T. marneffei.* The residual IL-1β response to *T. marneffei* in BMDCs from *Casp-1*^−/−^ and *Nlrp3*^−/−^ mice (Fig 4D), and particularly the lower fungal load and better survival in *Casp-1*^−/−^ mice compared to *Nlrp3*^−/−^ mice (Fig 7A-C), suggests possible contributions from alternate signaling pathways. For example, the engagement of Dectin-1 by *P. brasiliensis* activates caspase-8 which drives the transcriptional priming and post-translational processing of pro-IL-1β. Interestingly, the canonical caspase-1 inflammasome pathway normally restricts the noncanonical caspase-8 inflammasome activation, therefore in the context of caspase-1/11 deficiency, the lack of inhibition from caspase-1 enables caspase-8-dependent IL-1β processing (Ketelut-Carneiro *et al*, 2018). A recent study show that IL-1α release via caspase-11-mediated pyroptosis induced by *P. brasiliensis* reprograms Th17 lymphocytes to produce high levels of IL-17 and enhances effector mechanisms by innate cells (Ketelut-Carneiro *et al*, 2019). Further studies are required to investigate alternative molecular pathways in the induction of inflammasome activation against *T. marneffei* infection.

DCs from patients with HIV was previously shown to exhibit poor activation of NLRP3 inflammasome in response to PAMPs and DAMPs, which could be the consequence of increased basal expression and activation of NLRP3 induced by HIV in chronic infection leading to immune exhaustion (Pontillo *et al*, 2012), as well as the inhibitory effect on NLRP3 inflammasome activity by type I interferon which is commonly elevated in plasma of HIV-infected patients (Reis *et al*, 2019; Hardy *et al*, 2013). In response to LPS stimulation, PBMCs from HIV patients express relatively lower levels of activated caspase-1 (Ahmad *et al*, 2002), and IL-1β production in DCs fails to be upregulated by either LPS (Pontillo *et al*, 2012) or ATP which classically induces canonical activation of NLRP3 (Reis *et al,* 2019). It is possible that functional defect of NLRP3 could represent an additional contributory factor to increased susceptibility to *T. marneffei* infection in HIV-positive individuals. Further studies on functional evaluation of NLRP3 and caspase-1 activation in HIV-positive patients infected with *T. marneffei* will be required.

This study enhances our understanding of host defense mechanisms against *T. marneffei*. Knowledge about human immune response towards *T. marneffei* will have therapeutic implications in managing patients suffering from this fatal infection. Delineation of the roles of various cytokines towards protection against *T. marneffei* will provide important information to the treatment of patients who have secondary immunodeficiency resulting from the use of biologics for immunological disorders. For example, with the increasingly wide clinical use of IL-1 receptor antagonist (e.g. Anakinra, Rilonacept) or neutralization antibodies (Canakinumab) for patients with auto-inflammatory or auto-immune diseases, and therefore clinicians should be alerted to the susceptibility and signs of talaromycosis when treating patients who reside in endemic regions where there is increased risk of environmental exposure to *T. marneffei*. Finally, the study of functional cellular response towards *T. marneffei* may provide breakthroughs for the discovery of monogenic immune defects in patients with talaromycosis which is not otherwise explained. It is possible that genetic defects in molecules involved in Dectin-1/Syk signaling pathway, NLRP3 inflammasome activation and induction of Th1 and Th17 immune responses might predispose to *T. marneffei* infection.

To conclude, in this study, we first demonstrate that *T. marneffei* yeasts rather than conidia induce IL-1β response, which is differentially regulated in distinct cell types. Our findings show that Dectin-1/Syk signaling pathway mediates pro-IL-1β production upon fungal stimulation. *T. marneffei* yeasts also trigger the assembly of NLRP3-ASC-caspase-1 inflammasome to facilitate IL-1β maturation. The activated inflammasome partially enhances Th1 and Th17 immune responses against fungal infection. *Nlrp3*^−/−^ and *Casp1*^−/−^ mice are more susceptible to *T. marneffei* yeasts with higher fungal load as compared to WT mice. Collectively, our study highlights the importance of NLRP3 inflammasome in host defense against *T. marneffei* infection and sheds new light on the interaction between host and fungal cells.

## Materials and Methods

### Fungi

*T. marneffei and C. albicans* were isolated from patients by the Department of Microbiology, Queen Mary Hospital. *T. marneffei* conidia were cultured on Sabouraud Dextrose Agar (SDA) plates at 25°C for 7 days, and then collected in sterile water by wet cotton swabs, washed 3 times and re-suspended in sterile water, and finally filtered through 40μm nylon filters. To prepare *T. marneffei* yeasts, *T. marneffei* conidia were cultured on SDA plates at 37°C for 10 days, and then collected in sterile 0.1% PBST by wet cotton swabs, washed 3 times and re-suspended in RPMI 1640 with shaking for 24 hr at 37°C, and finally filtered through 40μm nylon filters. *C. albicans* were cultured on SDA plates at 30°C for 48 hr. Then, they were transferred to Sabouraud Dextrose (SD) broth and incubated at 37°C with shaking for 16–24 hr to obtain the yeast form of *C. albicans*. Simultaneously, *C. albicans* were cultured with shaking in enriched media consisting of RPMI+10% fetal bovine serum (FBS) at 37°C for 24 hr to obtain the pseudo-hyphae form of *C. albicans*. For fixed fungal preparation, fungi were washed in sterile PBS, fixed in 4% paraformaldehyde (PFA) solution for 10 min, and then washed three times in sterile PBS. For heat-killed fungal preparation, fungi were boiled at 100°C for 40 min.

### Cell isolation and culture

Peripheral blood mononuclear cells (PBMCs) were isolated from the buffy coats of healthy donors according to the principles of the Declaration of Helsinki and were approved by the responsible ethics committee (Institutional Review Boards of the University of Hong Kong/Hospital Authority Hong Kong West Cluster). Human primary CD14^+^ monocytes and CD4^+^ T cells were isolated from PBMCs by magnetic labeling with CD14 or CD4 microbeads respectively, followed by positive selection with MACS columns and separators (Miltenyi Biotec). For preparation of monocyte-derived macrophages, PBMCs were cultured in 10cm dish, and medium was changed gently at 2 hr after cell seeding. Cells were left overnight, and adherent cells were treated with cold 5mM EDTA for 10 min, scrapped and cultured in 24-well plates for 12 days. During this period, medium was changed on day 7 and day 9 to remove non-adherent cells. Human PBMCs, CD14^+^monocytes, monocyte-derived macrophages were cultured in RPMI 1640 supplemented with 5% heat-inactivated autologous sera and 1% penicillin/streptomycin. For monocyte-derived dendritic cells (DCs), CD14^+^ monocytes were cultured for 5 days in RPMI1640 containing 10% FBS, 1% penicillin/streptomycin, human IL-4 (40ng/ml, PeproTech) and GM-CSF (50ng/ml, PeproTech). Non-adherent or loosely adherent cells were collected.

Murine bone marrow-derived dendritic cells (BMDCs) were prepared as previously described (Helft *et al*, 2015). Briefly, bone marrow cells were isolated from the femurs and tibias of mice. Red blood cells were lysed on ice for 5 min by using 1x RBC Lysis Buffer (BioLegend, 420301). Bone marrow cells (3×10^6^ per well) were cultured in 6-well plates in 3 ml of complete medium (RPMI 1640, 1% penicillin/ streptomycin, 10% FBS, 1% Glutamax, 1% sodium pyruvate, and 55μM β-mercaptoethanol) supplemented with murine GM-CSF (20 ng/ml, Peprotech). Half of the medium was removed on day 2 and fresh medium supplemented with murine GM-CSF (40 ng/ml) was added. The culture medium was entirely aspirated on day 3 and replaced by fresh medium containing GM-CSF (20 ng/ml). Non-adherent cells in the culture supernatant and loosely adherent cells were harvested on day 6. The experiments that were performed for Western blot analysis of cell culture supernatant were carried out in Opti-MEM^TM^ serum free medium (Gibco). Murine splenocytes were prepared by homogenization of murine spleen, and subsequent lysis of red blood cells, and cultured in the complete medium.

### Cell stimulation

PBMCs (4×10^6^/ml) were co-cultured with live, PFA-treated or heat-killed *T. marneffei*, as well as live, heat-killed or PFA-treated *C. albicans* yeasts or pseudohyphae at 0.5 multiplicity of infection (MOI) for indicated time points. PBMCs (4×10^6^/ml), CD14^+^monocytes (2×10^6^/ml), monocyte-derived macrophages (1×10^6^/ml), or monocyte-derived DCs (1×10^6^/ml) were primed with LPS (10ng/ml for PBMCs and monocytes, 100ng/ml for monocyte-derived macrophages and monocyte-derived DCs, Invivogen) for 3 hr and then stimulated with heat-killed *T. marneffei* yeasts at 0.5 MOI for 18 hr. Human CD14^+^ monocytes (2×10^6^/ml) were pre-incubated with human anti-Dectin-1 (10μg/ml, Invivogen), anti-TLR2 blocking antibodies (10μg/ml, Invivogen), or corresponding isotype controls (10μg/ml, Invivogen), or different concentrations of R406 (Invivogen), or Z-YVAD(OMe)-FMK (ALX-260-074, Enzo) for 1 hr before co-culture with heat-killed *T. marneffei* yeasts for 18 hr. Murine BMDCs (2×10^6^/ml) were primed with or without LPS (100 ng/ml) for 3 hr, and then co-cultured with heat-killed *T. marneffei* yeasts at 1 MOI for additional 18 hr. Alternatively, Nigericin (10 μM) was added into LPS-primed cells in the last 40 min for positive control. Total CD4^+^ T cells (1×10^5^/well) were co-cultured for 7 days or 12 days with monocytes (1×10^5^/well)) or monocyte-derived DCs (0.2×10^5^/well) that were pre-pulsed for 3 hr with heat-killed *T. marneffei* yeasts (1×10^5^/well) or *C. albicans* yeasts (1×10^5^/well) in 96-well plates. Moreover, murine splenocytes (2×10^6^/ml) were co-cultured with heat-killed *T. marneffei* yeasts at 1 MOI for 0, 2, 6 day(s).

### Cytokine assays

Cytokine concentration in culture supernatant was measured by commercial ELISA kits: human IL-1β (ebioscience), human TNF-α, IL-18, and IFN-γ, IL-17A, IL-12p70, and IL-23 (R&D systems), and murine IL-1β, TNF-α, IFN-γ and IL-17A (R&D systems) according to manufacturer’s instructions. IL-6, MCP-1, IL-8, IL-18 and IFN-γ from human PBMCs were measured by LEGENDplex™ Human Inflammation Panel (Biolegend) according to manufacturer’s instructions.

### Western blot

Total cell lysates were prepared using sample buffer containing 5% β-mercaptoethanol. Protein concentration was determined by bicinchoninic acid (BCA) assay. Concentrated serum free culture supernatant was prepared as previously described (Hornung et al., 2008). Briefly, cell culture supernatant was precipitated by adding an equal volume of methanol and 0.25 volumes of chloroform. It was vigorously vortexed and centrifuged at 20,000 g for 10 min, and the upper phase was aspirated and discarded, and 500 μl methanol was added and centrifuged at 20,000 g for 10 min. Then protein pellet was dried in 55°C water baths for 5 min. Proteins from cell lysates (20-50 μg) or concentrated supernatant were boiled, separated on SDS-PAGE, and transferred onto PVDF membranes (Bio-Rad Laboratories). Membranes were blocked with 5% bovine serum albumin (BSA) in Tris-buffered saline (TBS) with 0.1% Tween 20 for 1 hr at room temperature, incubated overnight at 4°C with the following primary antibodies diluted 1:1000 in 5% BSA/TBST: anti-pro-IL-1β (#12703, Cell Signaling Technology) and anti-NLRP3 (AG-20B-0014-C100, Adipogen Life Sciences), and anti-mouse caspase-1 (AG-20B-0042-C100, Adipogen Life Sciences), and then incubated for 1 hr with goat anti-rabbit and goat anti-mouse secondary antibodies, respectively, at room temperature. Membranes were washed and exposed by adding enhanced chemiluminescence (ECL) in dark room. When necessary, stained membranes were incubated in stripping buffer at 65°C for 10 min, and washed 3 times with 0.1% TBS-Tween 20, and blocked with 5% BSA for 1 hr, and stained with anti-β-actin rabbit mAb (4970S, Cell Signaling Technology) followed by goat anti-rabbit secondary antibody staining and film exposure.

### Confocal imaging of ASC specks

Murine BMDCs (2×10^6^/ml) were seeded on the chamber slides (154941, Nalge Nunc International) and were co-cultured with heat-killed *T. marneffei* yeasts at 1 MOI for 18 hr, or were stimulated with LPS (100 ng/ml) for 18 hr with addition of Nigericin (10 μM) in the last 40 min. BMDCs were fixed with 4% PFA for 20 min and permeabilized with 0.1% Triton-X-100 for 10 min. Cells were washed 3 times and blocked with 5% BSA for 1 hr, followed by incubation with anti-ASC rabbit polyclonal antibody (SC-22514-R, Santa Cruz Biotechnology) overnight at 4°C. After washing, cells were incubated with AlexaFlour-488 conjugated goat anti-rabbit secondary antibody (Life Technologies) at room temperature for 1 hr. Subsequently, cells were stained with Alexa Fluor 647 phalloidin (ThermoFisher Scientific, A22287) for 30 min at room temperature avoiding light. Finally, slides with cells were mounted with ProLong Gold Antifade Mountant with DAPI (Invitrogen) and analyzed by confocal microscopy (Zeiss LSM 710). For quantification of ASC speck, 40-150 monocytes per field under microscopy with the magnification of 40x objective were manually analyzed, and the ratio of ASC speck positive cells to total cells was calculated. At least 3 fields per sample were counted.

### Flow cytometry

To detect Syk phosphorylation, CD14^+^ monocytes (2×10^6^/ml) were co-cultured with heat-killed *T. marneffei* yeasts at 0.5 MOI for 18 hr. Cells were fixed with BD Phosflow Fix Buffer I for 10 min at 37°C, followed by permeabilization with BD Phosflow Perm Buffer III for 30 min on ice. After washing, cells were stained with PE-conjugated phospho-Syk (Tyk525/526) antibody (#6485, Cell Signaling Technology) for 30 min on ice, and analyzed by flow cytometer (LSR II, BD). For intracellular staining of IFN-γ and IL-17A, CD4^+^ T cells were co-cultured with monocytes or monocyte-derived DCs in the presence of *T. marneffei* yeasts or *C. albicans* yeasts for 7 days or 12 days. Cells were then stimulated with PMA and ionomycin for 5 hr and Golgi plug (BD Biosicence) was added in the last 4 hr. Subsequently, cells were stained for 30 min at 4°C with FITC-conjugated anti-human CD3 antibody (Biolegend), followed by fixation and permeabilization of cells with BD Cytofix/Cytoperm™ (BD Bioscience). After washing, cells were stained with Alexa Fluor^®^ 647-conjugated anti-human IFN-γ antibody (502516, Biolegend) and PE-conjugated anti-human IL-17A antibody (512305, Biolegend) for 30 min at 4°C. Results were analyzed by flow cytometer (LSR II, BD).

### Murine systemic talaromycosis model

For *in vivo T. marneffei* yeasts infection, WT, *Nlrp3*^−/−^, *Casp1*^−/−^ mice were intravenously injected with 100 μl of suspension containing 5×10^5^ colony forming units (CFUs) live *T. marneffei* yeasts in sterile PBS. Mice survival was daily monitored following infection and mice were sacrificed once they showed signs of the humane endpoints. In the second batch of infection experiments, mice were sacrificed, and spleens and livers were harvested at 7 or 14 days post infection (dpi) of *T. marneffei* yeasts. Part of spleen and liver were weighed and homogenized in sterile PBS, and a series of diluted solutions of cell suspensions were plated onto SDA plates. Fungal load was assessed after culturing for 2 days at 25°C. Part of spleen and liver were fixed in 4% paraformaldehyde (PFA) and paraffin slides were prepared for hematoxylin and esosin (HE) staining and Grocott’s methenamine silver (GMS) staining. The stained sections were mounted and observed under microscope for histopathological assessment.

### Data analysis

Statistical analysis was carried out using GraphPad Prism v6.0 software. Figure 5G and H were ploted by R software, version 3.6.1. Data were represented as mean ± SEM. Statistical significance was determined by two-tailed Student’s t test, one-way ANOVA or two-way ANOVA with multiple comparison tests, log-rank test or Gehan-Breslow-Wilcoxon test for comparison of survival rate. p<0.05 was considered statistically significant.

## Acknowledgements

We gratefully acknowledge the Laboratory Animal Unit, the Faculty of Core Facility, and the Histopathological Service Center at the Department of Pathology, the University of Hong Kong, for their professional technical supports. This work was supported by the Edward and Yolanda Wong Fund, the RGC General Research Fund (GRF) (HKU project code: 17111814), and the Health and Medical Research Fund (No. HKM-15-M07 [commissioned project]).

## Author contributions

HM, JC, YL, PW, PL designed the experiments and analyzed the results. HM, YT, LK, CT, SP performed the experiments. HM and PL wrote and revised the manuscript.

---

**Figure EV1.**
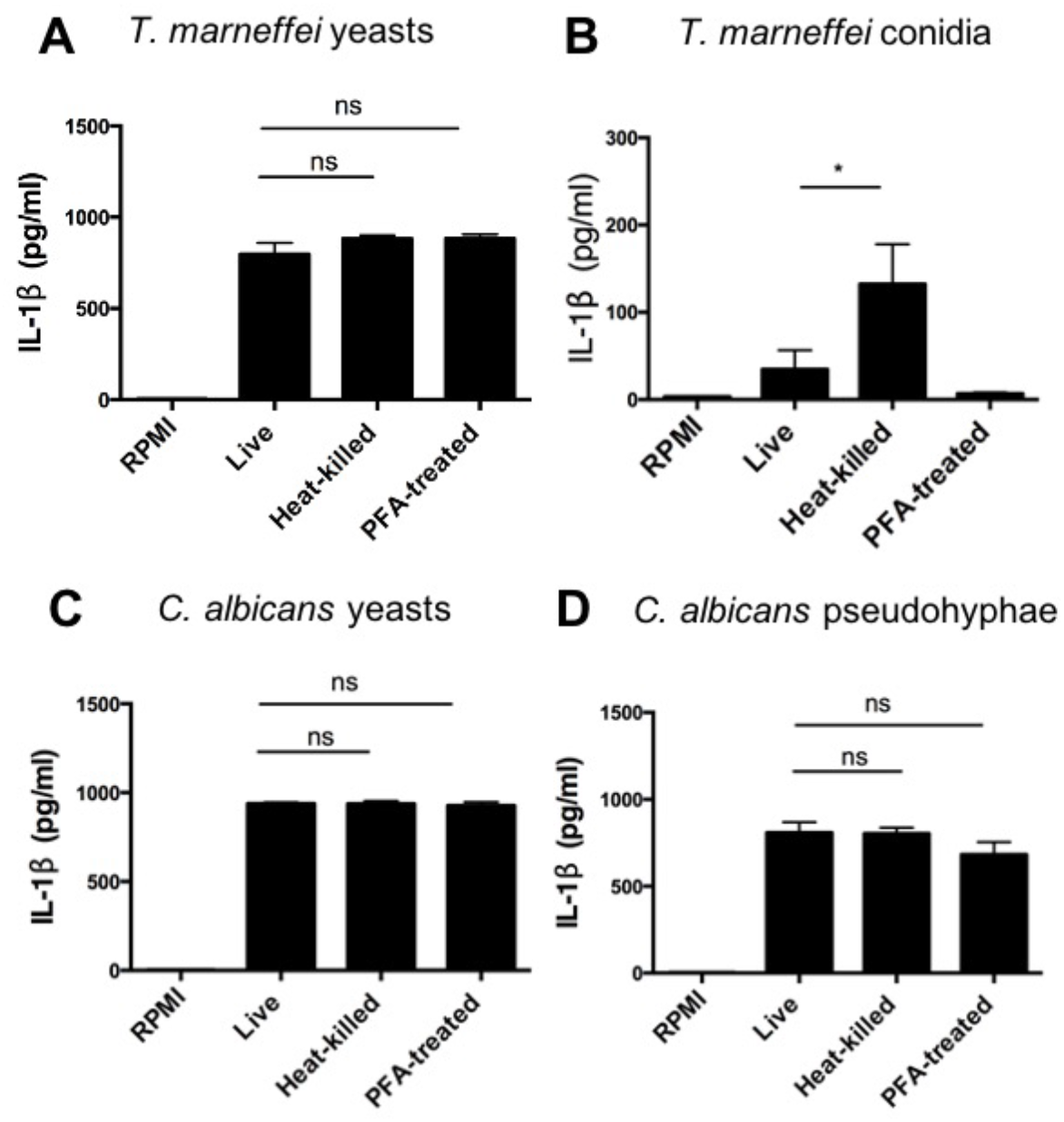
The viability of *T. marneffei* yeasts is dispensable for IL-1β response in human PBMCs. A, B. Quantification of IL-1β by ELISA in the cell supernatant of human PBMCs (4×10^6^/ml) stimulated with live, heat-killed or PFA-treated *T. marneffei* yeasts (0.5 MOI) and conidia (0.5MOI) (n=5) for 18 hr. C, D. Quantification of IL-1β by ELISA in the cell supernatant of human PBMCs (4×10^6^/ml) stimulated with live, heat-killed or PFA-treated *C. albicans* yeasts (0.5 MOI) and pseudohyphae (0.5MOI) (n=5) for 18 hr. Data information: In (A-D), data are depicted as mean ± SEM, and are analyzed by one-way ANOVA. ns =not significant, *p<0.05.

**Figure EV2.**
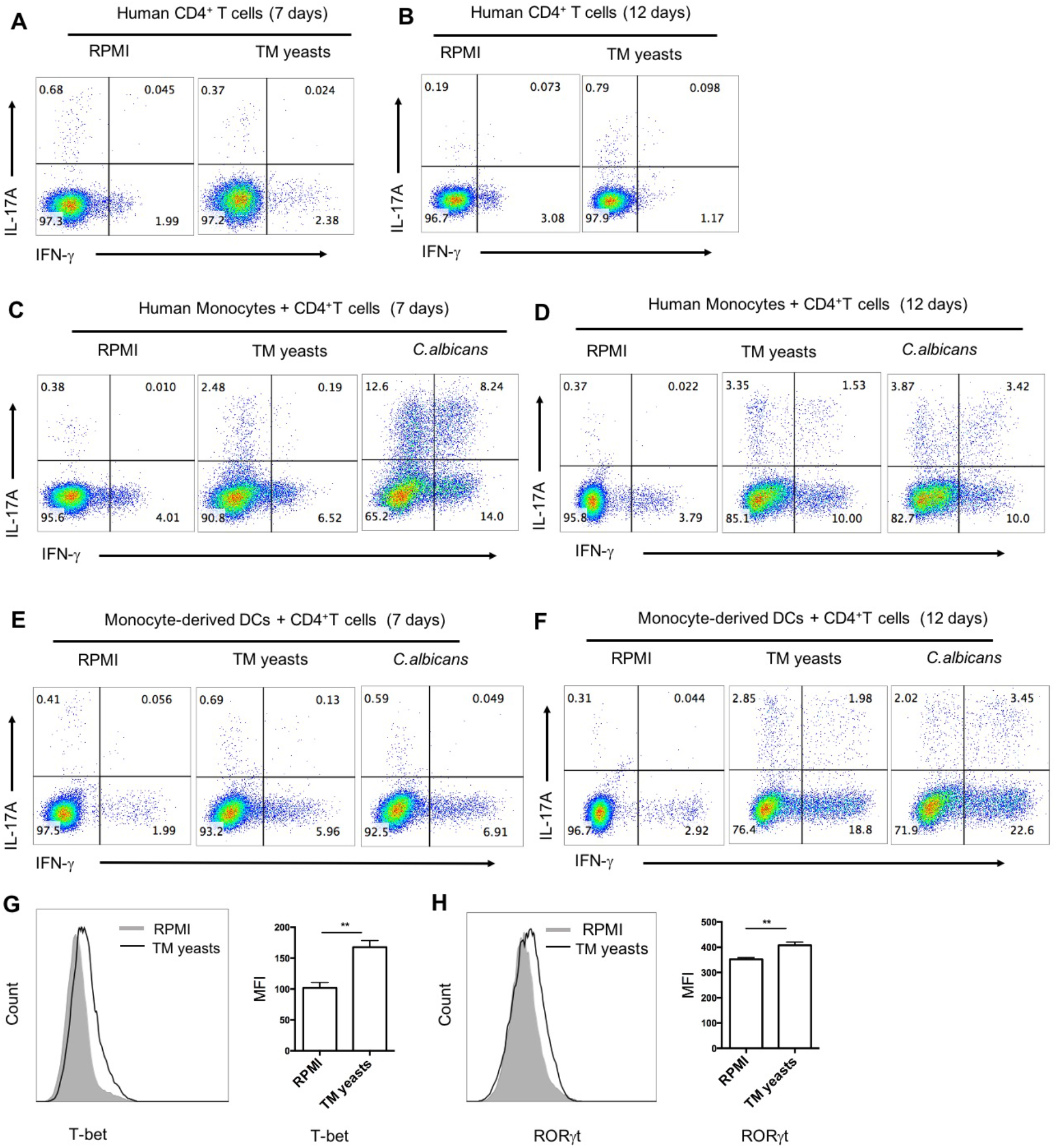
*T. marneffei* yeasts elicit differential Th1 and Th17 immune responses in human CD14^+^ monocytes and monocytes-derived DCs. A, B. Representative flow cytometric graphs of IFN-γ and/or IL-17A producing CD4^+^ T cells (1×10^5^/well) in human CD4^+^ T cells stimulated with heat-killed *T. marneffei* (TM) yeasts in the absence of autologous CD14^+^ monocytes or monocytes-derived DCs for 7 days (A) and 12 days (B). C, D. Representative flow cytometric plots of IFN-γ and/or IL-17A producing CD4^+^ T cells (1×10^5^/well) in human CD4^+^ T cells stimulated with heat-killed *T. marneffei* (TM) yeasts in the presence of autologous CD14^+^ monocytes (1×10^5^/well) for 7 days (C) and 12 days (D). E, F. Representative flow cytometric plots of IFN-γ and/or IL-17A producing CD4^+^ T cells in human CD4^+^ T cells (1×10^5^/well) stimulated with heat-killed *T. marneffei* (TM) yeasts in the presence of autologous monocytes-derived DCs (0.2×10^5^/well) for 7 days (E) and 12 days (F). G, H. Flow cytometric analyses of T-bet (G, n=5) and RORγt (H, n=4) in CD4^+^ T cells co-cultured with TM yeasts-primed autologous monocytes for 7 days. Data information: In (G and H), data are depicted as mean ± SEM, and are analyzed by two-tail Student’s t tests. “MFI” denotes “median fluorescence intensity”. **p<0.01.

**Figure EV3.**
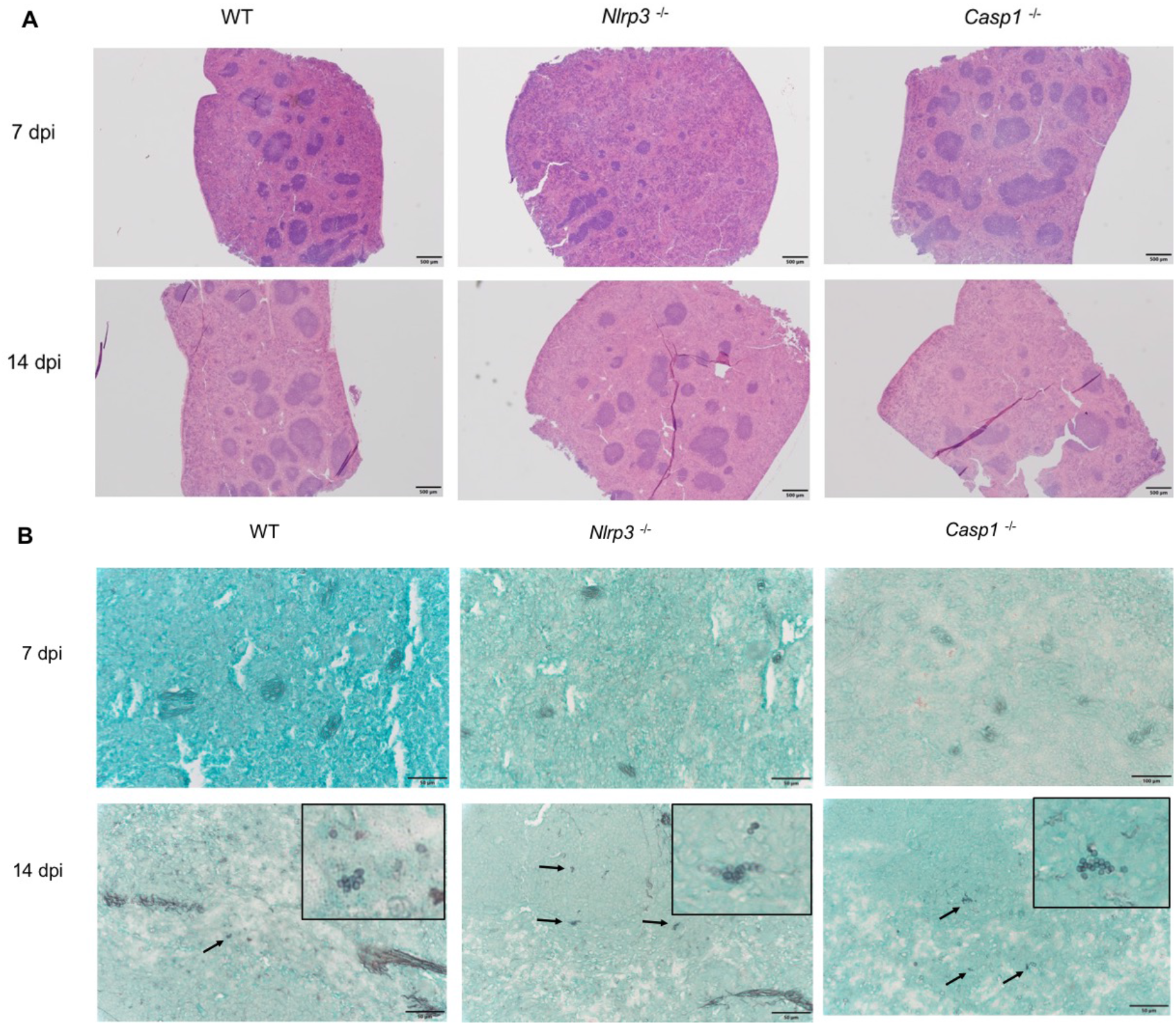
Histopathological analysis of murine livers and spleens. A. Representative graphs of HE staining of spleens from WT, *Nlrp3*^−/−^ and *Casp-1*^−/−^ mice intravenously infected *T. marneffei* yeasts (5×10^5^ CFU per mouse) at 7 days and 14 days post infection (dpi) (n=4). Scale bar denotes 500 μm. B. Representative graphs of GMS staining of spleens from WT, *Nlrp3*^−/−^ and *Casp-1*^−/−^ mice intravenously infected *T. marneffei* yeasts (5×10^5^ CFU per mouse) at 7 days and 14 days post infection (dpi) (n=4). Scale bar denotes 50 μm. Arrows indicate *T. marneffei* yeasts, and the insets indicate the magnified areas containing *T. marneffei* yeasts.

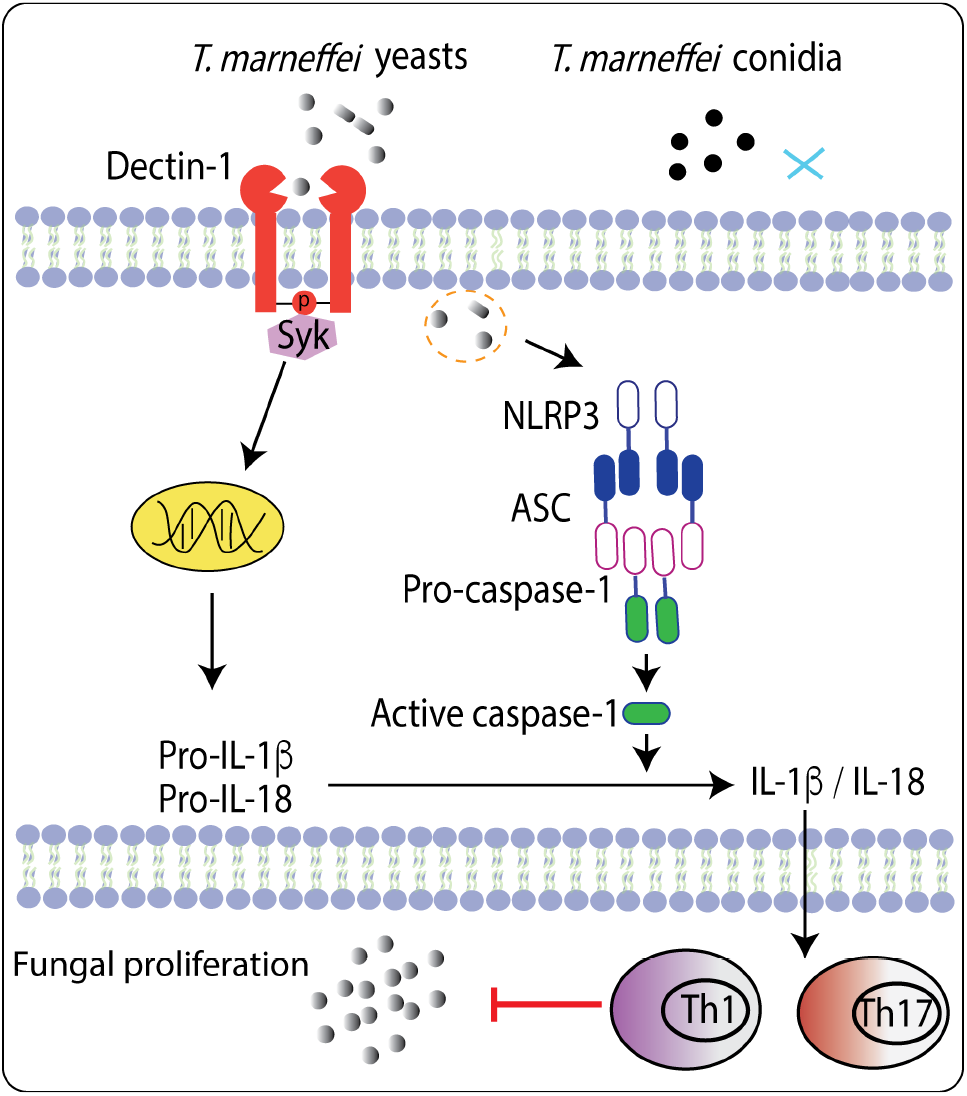
Graphical abstract.

